# Prioritizing transcription factor perturbations from single-cell transcriptomics

**DOI:** 10.1101/2022.06.27.497786

**Authors:** Rohit Singh, Joshua Shing Shun Li, Sudhir Gopal Tattikota, Yifang Liu, Jun Xu, Yanhui Hu, Norbert Perrimon, Bonnie Berger

**Author notes:** contributed equally (Rohit Singh, Joshua Shing Shun Li, Sudhir Gopal Tattikota).

## Abstract

The explosive growth of regulatory hypotheses from single-cell datasets demands accurate prioritization of hypotheses for *in vivo* validation. However, current computational methods emphasize overall accuracy in regulatory network reconstruction rather than prioritizing a limited set of causal transcription factors (TFs) that can be feasibly tested. We developed Haystack, a hybrid computational-biological algorithm that combines active learning and the concept of optimal transport theory to nominate and validate high-confidence causal hypotheses. Our novel approach efficiently identifies and prioritizes transient but causally-active TFs in cell lineages. We applied Haystack to single-cell observations, guiding efficient and cost-effective *in vivo* validations that reveal causal mechanisms of cell differentiation in *Drosophila* gut and blood lineages. Notably, all the TFs shortlisted for the final, imaging-based assays were validated as drivers of differentiation. Haystack’s hypothesis-prioritization approach will be crucial for validating concrete discoveries from the increasingly vast collection of low-confidence hypotheses from single-cell transcriptomics.

## Introduction

Identifying the causal regulatory transcription factors (TFs) that control cell differentiation is a fundamental question in biology. Since causality can only be hypothesized, but not confirmed from observational studies^1^, determining a causal role for any TF requires extensive perturbatory validations *in vivo*. However, such perturbations are time-consuming and expensive. Consider a relatively simple scenario where one needs to analyze 10 homozygous mice that can only be generated from breeding heterozygous animals. A total of 40 offspring would be required to obtain these 10 (25%) homozygotes, imposing about 6 months of delay and incurring thousands in costs (**Supplementary Note 1**). Therefore, being able to prioritize causal TFs likely to have a causal role has long been an important consideration. In this paper, we focus on prioritization of perturbations likely to have a high success rate, enabling more efficient experimentation. Traditionally, in situ and immunohistochemistry approaches have been used to direct downstream perturbation efforts to shortlist putatively causal TFs, with the general intuition being that lineage-determining TFs should be active primarily in undifferentiated cells.

Single-cell RNA-sequencing (scRNA-seq) has emerged as a powerful alternative to these traditional approaches for generating causal regulatory hypotheses. Since scRNA-seq can capture gene expression profiles that span the differentiation landscape, it should be possible to identify and prioritize likely-causal TFs from this data. However, existing computational approaches do not directly address this challenge. These approaches often optimize metrics like area under receiver operating characteristic curve (AUROC) that are broad and assign substantial importance to low-confidence inferences. Unfortunately, this may come at the expense of precision in high-confidence predictions, wasting valuable experimental time and resources. Current inference approaches can be broadly categorized into two groups. The first category consists of methods aimed at robustly determining whether a TF is active in a cell. This is non-trivial, since the raw transcript counts of a TF may not be an accurate representation of its activity. Thus, methods like SCENIC^2^, SCENIC(+)^3^ and CellOracle^4^ incorporate other data modalities (e.g., the co-expression of TF target genes) for greater robustness. The second category of methods estimates the cell differentiation course (i.e., pseudotime) and infers regulatory relationships consistent with the estimated pseudotemporal ordering. Prioritization of lineage-determining TFs requires accuracy in assessing whether, and where, the TF is active. On its own, neither of the above two approaches is sufficient: multimodal approaches (like SCENIC) do not account for the pseudotime landscape while pseudotime-based gene regulatory network (GRN) inference methods are vulnerable to the noise and sparsity of scRNA-seq data^5^. Integration of the two approaches could address this challenge but off-the-shelf techniques yield suboptimal results. For instance, some studies calculate the differential estimates of TF activity between discrete cell clusters which discretizes the differentiation landscape. An alternative would be to correlate or regress a per-cell TF activity measure (e.g., from SCENIC) with the cell’s pseudotime score. This can handle continuous differentiation landscapes but can struggle with non-uniform distributions of cells across pseudotime. Since pseudotime scores emerge from graph-theoretic analyses of the single-cell landscape, there is no guarantee that cells will be uniformly distributed across the score range (**Figure 2I,J**). For these reasons, we sought a method that could effectively combine these broad approaches, as part of a cohesive scheme to prioritize regulatory hypotheses.

We introduce Haystack, a hypothesis prioritization-validation protocol for efficiently identifying causal TFs in the cell differentiation landscape. Our approach achieves greater confidence in its top-ranked TF candidates by integrating the two aforementioned inference strategies, rather than seeking to improve the approaches individually. Having a high hit-rate in the top predictions ensures that only a few TFs need to be validated in vivo, increasing experimental efficiency. Starting from a scRNA-seq study of differentiating tissue(s), we ultimately validate one or more TFs whose *in vivo* perturbation results in phenotypic changes in cell lineages. Our design goal was to achieve a high hit-rate in biological validations, especially during the final microscopy-based imaging or other confirmatory analyses. In Haystack, we introduce two key conceptual advances. The first is applying the principles of active learning, a subfield of machine learning, where an iterative approach is used to refine a set of hypotheses. To do so, Haystack switches between computational and biological modes where the results of intermediate biological experiments are computationally analyzed to hone in on a smaller set of higher-confidence candidates for detailed validation. The second key advance is achieving high early-precision in the initial computational prioritization of TFs by leveraging the concept of optimal transport (OT). We combine multimodal and pseudotime-based inference methods by first applying multimodal techniques to robustly estimate TF activity. The first phase of Haystack is to map each TF’s activity as a probability distribution over pseudotime and apply the OT metric to prioritize TFs with activity concentrated in specific regions of the landscape. We do so by computing the OT cost of transforming a TF’s probability distribution into an uninformative, uniform activation over all cells. This OT score is augmented with a feature selection score to further improve the precision of the top predictions. This set of high-prioritized TFs is further refined with another OT computation to hone in on “source” TFs that have likely downstream targets, generating a small set of causal high-confidence TF hypotheses. As a preliminary evaluation, we applied our OT algorithm on scRNA-seq studies of the mouse gut and human leukemia. Over 75% of the TFs prioritized by Haystack had evidence from the literature supporting a causal role in differentiation. The second phase of Haystack’s active learning protocol involves exploratory quantitative real time PCR (qRT-PCR) that is applied to further prioritize one or two TFs for final validation. With all of these steps together, Haystack enabled efficient *in vivo* validations of regulatory hypotheses prioritized from *Drosophila* blood and gut scRNA-seq datasets^6,7^. As a result, we efficiently identified novel TFs involved in the differentiation of both tissues. Investigating the poorly studied lamellocyte (LM) differentiation trajectory, our analysis revealed the TFs *Xbp1* and *CG3328* as playing a causal role in the differentiation of LMs. In gut tissues, Haystack revealed that the TF *pebble (peb)* helps drive the differentiation of intestinal stem cells into enterocytes, resolving two previous studies that showed opposing results. Altogether, Haystack enables prioritization of a small set of lineage determining TFs that could be easily tested *in vivo*, thereby limiting laborious validations by excluding low-confidence hypotheses from scRNA-seq datasets.

## Results

### Design of Haystack: prioritizing TFs using optimal transport

Haystack is motivated by the biological intuition that a lineage-determining regulator should be active primarily in undifferentiated cells, and should have as targets transcriptional modules active in terminally differentiated cells (**Figure 1**). The computational challenge we address is of implementing this intuition to achieve high precision (i.e., score hypotheses such that the top-ranked ones have a higher hit rate) while retaining robustness to scRNA-seq data noise and sparsity.

**Figure 1.**
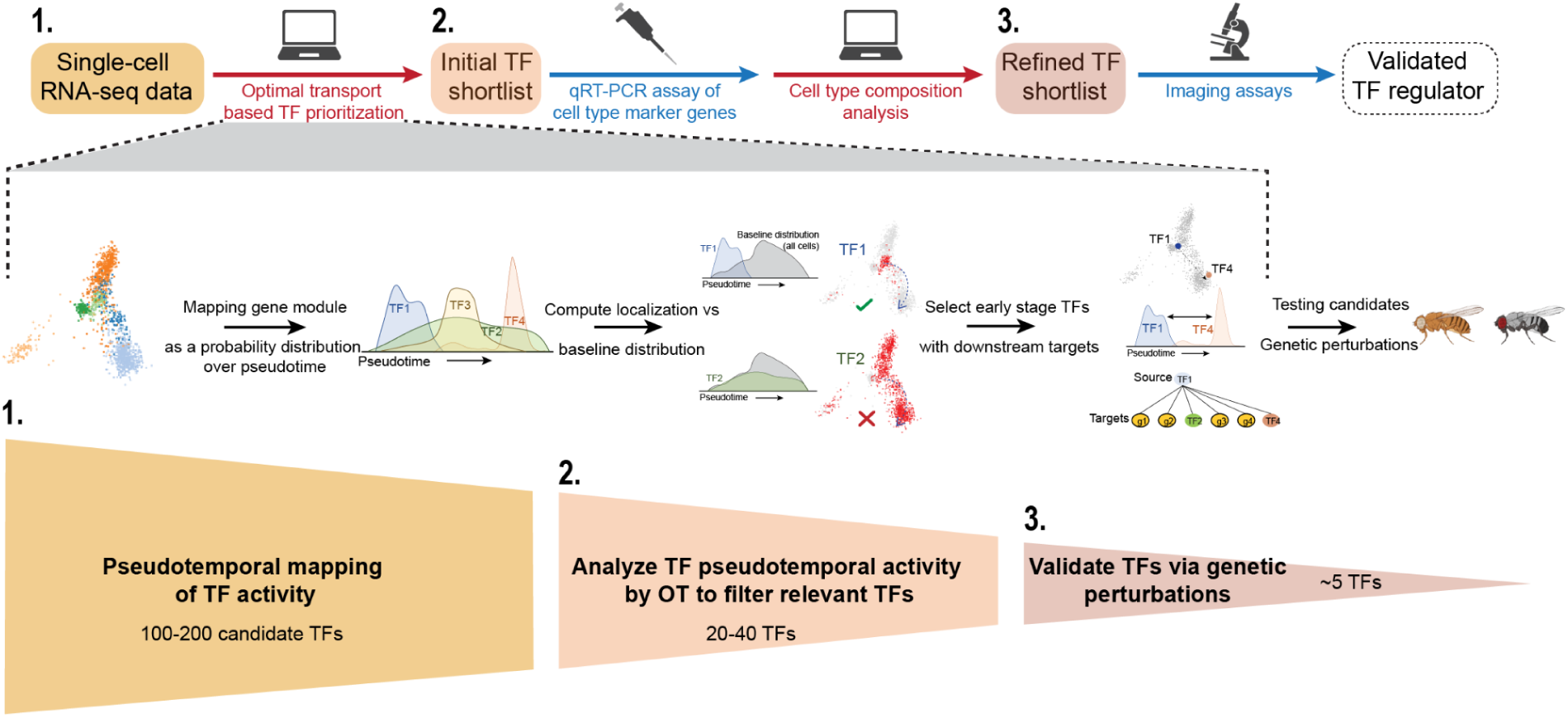
Prioritization-validation of causal regulatory hypotheses: Haystack workflow. Haystack’s hybrid computational-experimental workflow prioritizes causal transcription factors (TFs) along a trajectory. Our goal is to shortlist a few high-confidence top hits so that *in vivo* perturbation efforts are not wasted. We combine robust inference of TF activity with trajectory inference, relying on the concepts of active learning and optimal transport. The key steps are numbered in the schematic. (**1**) Starting from an observational scRNA-seq study of differentiating or developing tissue, we computationally reconstruct the differentiation landscape (e.g., by pseudotime). (**2**) Applying optimal transport theory, we combine robust estimates of TF activity that also consider a TF’s targets (regulons) with pseudotime information for an initial prioritization of TFs. (**3**) Cell-type composition inferred from qRT-PCR experiments further selects TFs whose perturbations result in changes in differentiation. Focused imaging-based validations are subsequently conducted on the highest-confidence candidates.

In Haystack, we model each TF activity as a probability distribution over cells, enabling us to leverage the principle of optimal transport (OT) to quantify the localization of a TF over the differentiation landscape (**Methods**). OT is a mathematical formulation for measuring the distance between two probability distributions under some cost function (**Figure 1**). The non-parametric nature of OT allows it to robustly handle varied differentiation landscapes. For instance, in a study of *Drosophila* gut differentiation, we observed pseudotime regions with low cell count (**Figure 2I**). Our OT based approach prioritized *EcR* regulon (which has been implicated in stem-cell differentiation in fly gut ^8^) while a correlation-based measure was led astray by abnormally low *EcR*-regulon readings in an early part of the differentiation regime with few cells (pseudotime scores 2-5, **Figure 2K**). We constrained Haystack to the processing of linear or diverging manifolds, which comprise the vast majority of cell differentiation scenarios.

**Figure 2.**
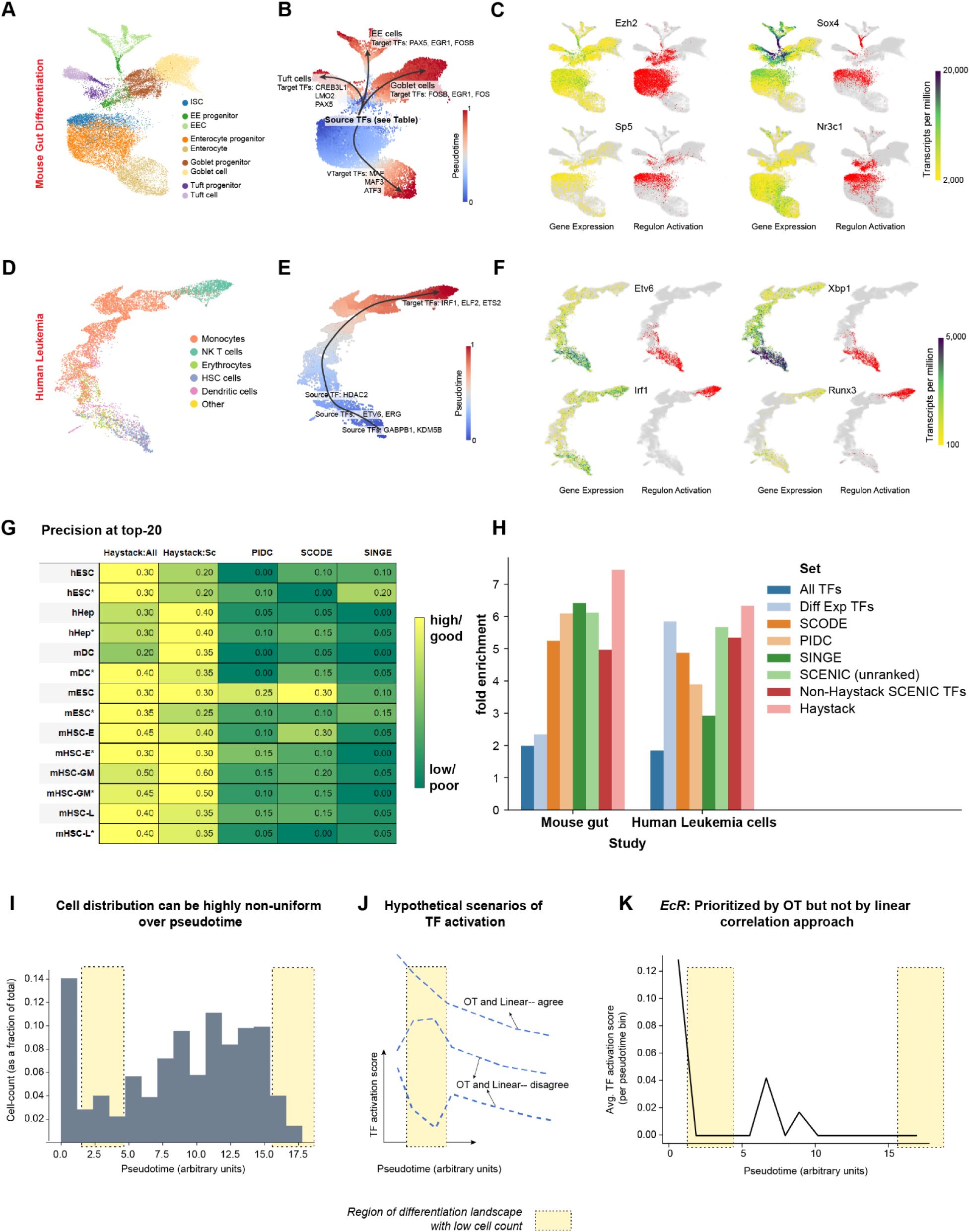
Haystack identifies biologically relevant TFs in mouse gut and human leukemia. **A-C.** (**Mouse gut**): Uniform Manifold Approximation and Projection (UMAP) plots depicting the re-clustering analysis of (**A**) scRNA-seq data of mouse gut (Bottcher et al., 2021); (**B**) Monocle 3-based pseudotime analysis reveals that ISCs can give rise to four lineages: Goblet cells, EECs, Tuft cells, and Enterocytes; and (**C**) gene expression (yellow-green color scale) of the TFs *Ezh2*, *Sox4*, *Sp5* and *Nr3c1*, contrasted with their TF-module activations (in red, inferred by SCENIC) in the respective intestinal clusters. **D-F.** (**Human leukemia**): Re-clustering of leukemia scRNA-seq data^11^ (**D**) identifies known blood cell populations; (**E**) Monocle 3-based pseudotime analysis reveals a single lineage trajectory with HSCs as the source; and (**F**) UMAP plots showing the expression (yellow-green color scale) of the TFs *Etv6*, *Xbp1*, *Irf1*, and *Runx1* compared to their TF-module activity (red).T **G-H.** (**Benchmarking**): The tabular plot (**G**) presents the precision of the top-20 predictions of Haystack (All: source and target TFs, Sc: source-only TFs), PIDC, SCODE and SINGE on a variety of mouse and human cell types (**Supplementary information** for details). The yellow–green color gradient in each row is scaled to ensure a uniform maximum (yellow) across all rows. The ground-truth gene sets are sourced from ChIP-seq data, with both cell-type specific and non-specific ChIP-seq data (the latter are indicated by an asterisk). (**H)** The bar graph represents enrichment scores of TF predictions against ground-truth gene sets obtained by literature text mining. The top 30 TFs (or fewer, as available) predictions derived from each GRN inference method are used to determine if they are enriched in a curated collection of Pubmed studies. The results are displayed as a fold-enrichment compared to the control of 30 random genes. As additional controls, enrichment scores when using all TFs (i.e., more than 30) or TFs differentially expressed between the initial and final cell types are also shown. **I-K**. (**Advantages of optimal transport formulation**): The histogram (**I**) represents the typical non-uniform distribution of cell counts (as a fraction of total cells) within different pseudotime intervals (here, *Drosophila* midgut differentiation). Yellow regions demarcate areas of pseudotime landscape with low cell counts. (**J**) A graph showing hypothetical scenarios of various TF activations as a function of pseudotime. Approaches that simply correlate (or linearly regress) TF activation against pseudotime can be confounded by anomalous readings in a few cells. In a simple case (top), where the TF profile is consistent across pseudotime regimes, the OT and correlation-based inferences agree on selecting the TF. However, low cell-count regions (bottom two profiles) of the differentiation landscape may misdirect the correlation-based approach when prioritizing top hits, while our OT-based approach that non-parametrically compares a TF’s activity profile to the background distribution of all cells is more effective. (**K)** An example, from our study of *Drosophila* midgut differentiation, where the OT-based approach detects a potentially interesting TF (*EcR*, known to be implicated in intestinal stem cell differentiation^8^) with early activity in the pseudotime landscape. In contrast, correlation produced a lower score, leading to a failure to select this TF.

Haystack utilizes SCENIC regulons ^2^ as a measure of TF activity at the level of modules. Each regulon relates to the activity of one TF, quantified not just in terms of its own gene expression but also that of its putative targets (as inferred from the cisTarget database of TF binding sites). After identifying the TF modules active in each cell, we characterize the localization of each TF along the differentiation time-course, seeking to hone in on TFs that are active in just one differentiation stage and not broadly active. To further refine the set of candidate TFs, we prioritize TFs more likely to play a causal role in differentiation. Lineage-determining master regulators typically have more than one downstream target^9^, with their ability to influence a broad transcriptional program being key to fate determination. Accordingly, we queried the cisTarget database to identify TF pairs (TF_source_–TF_target_) where the binding site of TF_source_ was found upstream of TF_target_, suggesting that TF_source_ might regulate TF_target_. From the shortlist of well-localized TFs, we considered every pairwise combination of TFs (say, TF_1_ and TF_2_) such that TF_1_ is active earlier in the pseudotime landscape than TF_2_ and with the binding site of TF_1_ upstream of TF_2_. We then applied OT to compute the pseudotime distance between the two TFs. From these pairs, we extracted the subset corresponding to high OT distance (**Methods for details**). The source TFs in these pairs are the candidate TFs we generate as the initial set of high-confidence hypotheses prioritized for experimental validation. Within this set, we rank TFs by the number of well-separated pairs in which the TF occurs as a source.

### Haystack prioritization algorithm recovers known regulators of mouse gut differentiation and human hematopoiesis

To assess the predictive potential of our prioritization algorithm, we applied Haystack on scRNA-seq studies pertaining to mouse gut differentiation (GSE152325^10^) and Acute Myeloid Leukemia (AML) cells (10.5281/zenodo.3345981^11^). In the mouse gut, intestinal stem cells (ISC) differentiate along four lineages: enteroendocrine (EE) cells, tuft cells, goblet cells and enterocytes (**Figure 2A,B**). We applied Haystack on each of the lineages, identifying the source TFs localized to or near the ISC cell type. Interestingly, when we visualized individual TF using transcript levels alone (eg. *Ezh2*, *Sox4*, *Sp5* and *Nr3c1*), a diffuse expression pattern was observed (**Figure 2C**). However, visualizing with TF modules revealed that the same diffusely expressed TFs had very localized module-level activity (**Figure 2C**). We aggregated these results across the lineages by counting the total number of downstream targets per source TF and ranked them within each lineage. Of the 24 putative source TFs predicted by Haystack, 75% had substantial support in the literature for their involvement in the development of the gut (**Table 1**) most of which were related to differentiation. For example, *Sox4* activity is localized to the intestinal progenitors and was previously shown to promote gut secretory differentiation towards tuft and EE fates^12^ (**Figure 2B, Table 1**). Another example is *Pou2f3*, where mice null for this TF lack tuft cells and also become defective in their response to helminth parasites^13^. Since the activity of *Sp5* is localized to ISCs, genetic perturbations might unravel new functions of *Sp5* previously unknown to ISC differentiation.

Next, we applied our prioritization algorithm to investigate hematopoiesis in human AML^11^. Pseudotime analysis of the data revealed that hematopoietic stem cells (HSC) differentiate into dendritic cells, erythrocytes, NK T cells and monocytes in a linear trajectory (**Figure 2D,E**). We again found that TF modules exhibited clearly localized activity while the raw gene expression was diffusely expressed (e.g., *Etv6*, *Xbp1*, *Irf1* and *Runx3,* **Figure 2F**). In total, we identified a total of 25 high confidence TF predictions, 85% of which had substantial evidence in the literature to support involvement in blood function. For instance, the TFs *GABPB1* and *KDM5B* that are localized at the HSC cluster (**Figure 2E**) are known to be involved in the maintenance and self-renewal potential of HSCs. Specifically, whilst *GABPB1* is required for stem/progenitor cell maintenance and myeloid differentiation in humans^14^, *KDM5B* is required for HSC self-renewal in mice and is specifically enriched in the human HSC compartment. Interestingly, both these TFs are implicated in leukemia demonstrating the power of our prioritization method in efficiently identifying TFs that are relevant in the context of hematopoiesis.

For a more systematic evaluation, we assessed Haystack within an existing framework for benchmarking GRN inference methods, based on ground-truth ChIP-Seq datasets across five cell types curated by Pratapa et al^5^. We compared Haystack prioritization against the top-ranking methods from that study. While standard machine learning metrics like area under precision recall curve give credit to an algorithm even for its weak predictions, we sought to focus our measurement on an algorithm’s top predictions, since those would be a biologist’s focus. Accordingly, we calculated early precision, a metric also considered by Pratapa et al. Compared to existing methods, Haystack achieved substantially higher precision on its top 10, 20 or 30 predictions, robustly outperforming them across different hyperparameter settings (**Figure 2G, S1A-F**). To assess our predictions for mouse gut development and human blood cell differentiation studies, we needed corresponding ground-truth gene sets. Pre-curated genes sets for these tissues being unavailable, we curated our own. For a given tissue, a collection of studies was first outlined as a testing set. This was defined by text-mining the Pubmed database, querying the titles and abstracts for words (**Methods**) that correspond to the tissue of our interest. The frequency at which a random set of 30 genes would appear in the testing set of studies was set as the baseline. Subsequent predictions were calculated as a fold-change in enrichment compared to this control. For example, simply selecting 30 TFs randomly led to a higher enrichment than control, as would be expected. We computed the enrichment score for the top-30 TF predictions generated from methods including SCODE^15^, PIDC^16^, SINGE^17^, Haystack. All methods had a significantly higher enrichment score when compared to the control. While some differences in the enrichment score were observed between the mouse gut and human leukemia, Haystack consistently scored highest (**Figure 2H**). Altogether, our hypothesis prioritization algorithm can efficiently recover known biology from scRNA-seq datasets and is a significant improvement to the predictive ability of the existing simple utilizations of GRN inference methods.

### Prioritization-validation of TFs along cell lineage trajectories in *Drosophila*

The second phase of Haystack consists of an initial qRT-PCR-based validation screen which is computationally analyzed to further prioritize the highest-confidence hits that are finally validated by a low-throughput imaging-based assay. Specifically, the OT-based shortlist of TFs generated from the first phase is investigated with qRT-PCR of cell-type markers, with changes in cell-type composition estimated by a novel aggregate-fold-change metric (**Methods**). TFs with supporting evidence are then assayed with microscopy (**Figure 1**).

As a proof of principle we employed the entire Haystack pipeline on *Drosophila* where putative predictions can be readily and cost-effectively tested with genetic perturbations. We focused on the adult fly midgut and larval blood for several reasons^6,7,18^: i) well-established tools are available to conduct genetic perturbations in a spatial and temporal manner; ii) the directionality of differentiation is relatively simple with limited intermediate-or terminal-states; iii) outlining TF-TF source-to-target associations can unravel TF cascades that underlie cell lineage progression in the two lineages.

### Initial validation using qRT-PCR based identification of cell type composition

Iterative experimentation is a key precept in active learning, the subfield of machine learning focused on optimal experiment design: initial experiments are designed to prune the search space or explore under-sampled regions, guiding future experimentation. Here, we apply this concept by leveraging qRT-PCR based intermediate validation to prune the set of prioritized TFs for more extensive downstream validation. Compared to the latter, which involves low-throughput phenotypic and imaging assays, qRT-PCR-based validation provides an accurate medium-throughput validation that is more efficient than imaging and phenotypic studies and allows multiple perturbations to be evaluated simultaneously.

From the original scRNA-seq study, we identified the cell types/clusters of interest and applied differential expression analysis to identify a limited set of markers (typically, 1-3) per cell type. In our *Drosophila* assays, this resulted in less than 15 markers in total, thus amenable to a single qRT-PCR study. We reasoned that concurrently assaying multiple markers per cell-type, made feasible by qRT-PCR, would enhance our ability to discern cell-types, providing an advantage over other approaches (e.g., immunohistochemistry) where only one marker per cell-type is used.

For each of the shortlisted TFs, we perform tissue-specific perturbation experiments and assayed cell-type markers to assess changes in cell-type composition as a result of the perturbation. However, a challenge when estimating cell-type composition changes using qRT-PCR is deconfounding tissue-level proliferation changes (which are not our primary interest) from differentiation changes (which are). Since qRT-PCR CT values are typically normalized against the background CT value of a housekeeping or ribosomal gene, this can confound cell type composition analysis. Accordingly, we introduce a fold-change metric to adjust for this confounding factor and robustly recover estimates of cell type compositions (**Methods**). For the final validation, we prioritized only the TFs where both overexpression and knockdown showed mutually consistent cell-type composition changes. Notably, every TF that passed the intermediate qRT-PCR validation test was also fully validated in downstream imaging studies, suggesting that Haystack prioritization process is broadly applicable across organisms and can successfully honed in on the TFs most likely to play a causal role.

### Applying Haystack prioritization-validation on adult midgut

The fly midgut consists of a monolayer of absorptive enterocytes (ECs) and secretory enteroendocrine cells (EEs) that are replenished by self-renewable intestinal stem cells (ISCs)^19,20^. Previously, we applied scRNA-seq on adult fly guts and identified a total of 22 clusters mainly consisting of finer sub-classifications of EEs and ECs^7,18^. Some of these subtypes are distinguished by their spatial location whereas others were intermediate states between ISC/EB and a specific terminal state^18^. Following the original study, we used SlingShot^21^ to map individual cells onto a pseudotime trajectory consisting of three lineages with the starting point set as ISC/EB (in blue) and end points as EE, aEC or pEC (red) (**Figure 3A-B**). By mapping TF module activity along these trajectories, Haystack prioritized eight TFs that were localized to the ISC/EB starting point (Source TFs). These included *Forkhead box K (FoxK)*, *cap-n-collar (cnc)*, *pebbled (peb)*, *P-element somatic inhibitor (Psi)*, *drumstick (drm)*, *TATA binding protein (Tbp)*, *Ten-Eleven Translocation family protein (Tet)* and *Mondo* (**Figure 3C**). Four out of the eight predictions (*FoxK*, *cnc*, *peb* and *drm*) were previously shown to play a role in the gut, highlighting the sensitivity of our prioritization algorithm^22–26^.

**Figure 3.**
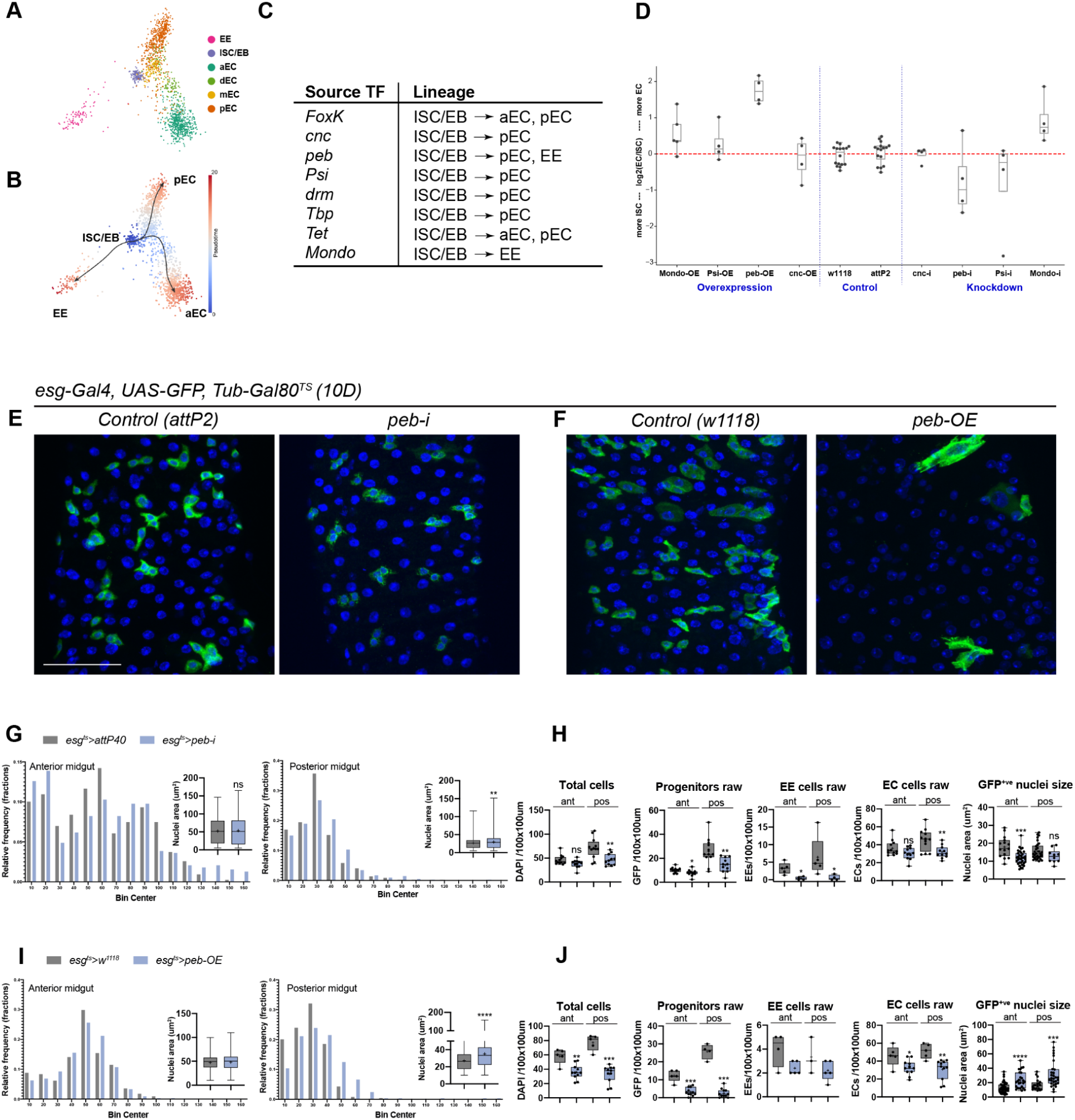
Analysis of Drosophila midgut scRNA-seq data by Haystack identifies known and novel TFs in the gut lineages. **A.** *Drosophila* midgut scRNA-seq dataset^7^ shows six known intestinal cell populations: enteroendocrine (EE), intestinal stem cells/enteroblasts (ISC/EB), enterocytes (ECs) of the different regions of the gut [anterior (aEC), differentiating (dEC), middle (mEC), and posterior (pEC)]. **B.** Pseudotime analysis using Slingshot reveals three distinct lineages from the ISC/EB cluster: EE, aEC, and pEC. **C.** Haystack identifies source TFs specific to three lineages. **D.** qRT-PCR-based lineage analysis of ISC and EC cell-types in the midguts of *Mondo*, *Psi*, *Peb*, and *cnc* perturbations compared to their respective controls (*w^1118^* and *attp2*). Only *Peb* displays mutually consistent overexpression and knockdown phenotypes. **E-F.** Confocal images of *Drosophila* midguts of *peb-i* and *peb-OE* show a decrease in progenitors. Scale bar = 100 um. **G-J.** Quantification of confocal micrographs after *peb* perturbation. **(G,H)** *peb* knockdown and **(I,J)** *peb* overexpression in intestinal progenitors. Parametric t-tests were used to calculate statistics for nuclei area. Non-parametric t-tests were used to calculate the statistics where *, **, ***, **** represent p values <0.05, 0.01, 0.001, 0.0001, respectively.

Next, we designed *in vivo* perturbation experiments to test whether these candidate TFs play a causal role in regulating ISC differentiation. Using differential gene analysis, we selected a total of 11 markers that were distinctly expressed in ISC/EB (*Dtg, N, LanB1*), aEC (*CG6295, Npc2f*), pEC (*LManVI, Mur29B*) or EE (*AstC, IA-2*) (**Figure S2A**). In the original Hung et al. study, these markers had been identified for individual cell clusters using a differential expression analysis in Seurat ^27^ and for each cell type, we chose a combination of markers such that all sub-clusters of a cell type were covered. We knocked down or overexpressed the eight TFs specifically in adult ISC/EB with publically available reagents. Among these, we found five RNAi lines (*peb-i, Psi-i, Tet-i, Mondo-i, cnc-i*) and seven overexpression (OE) lines (*peb-OE, Psi-OE, FoxK-OE, drm-OE, two Mondo-OE, cnc-OE*) that successfully reduced or increased, respectively, the gene of perturbation when assaying mRNA extracted from the whole midgut (**Figure 3C, S2**). We measured the levels of the 11 markers for each perturbation, which amounted to a total of 132 observations (Figure S2). We calculated the fold change (FC_RpL32_) in comparison to a control, using RpL32 as a reference. In all cases, perturbation led to at least one, and up to 11, significant fold change(s) in marker gene expression. The FC_RpL32_ in markers gave us a proxy for the cell-type composition of the gut. For example, knockdown of *Psi* in intestinal progenitors caused a decrease in EE and pEC markers whilst ISC/EB and aEC markers remained unchanged (except for *Npc2f*). This suggests that *Psi* is involved in promoting the differentiation of progenitors to EEs and ECs in the posterior midgut (**Figure 3D**). *Drm* overexpression caused an increase in ISC/EB markers with no changes in most terminal cell-type markers other than an increase in *Mur29B*, suggesting that *drm* promotes ISC proliferation but is not involved in differentiation (**Figure S2**). To systematize these comparisons, we applied the fold-change metric described above.

We focused on genes (*Psi*, *peb*, *cnc* and *Mondo*) where we had both RNAi and OE to confirm opposing effects of both perturbations. We used the EC–ISC mean marker ratio to compare our perturbations with the control. The most striking difference we observed was with *peb* perturbations. *Peb* OE increased EC, whereas *peb* knockdown decreased the EC–ISC ratio (**Figure 3D**). Interestingly, two independent groups had previously identified *peb* to function in the fly midgut ^22–26^. However, their findings were in conflict: Zeng et al. found that the knockdown of *peb* caused ISC-to-EC differentiation. In contrast, Baechler et al. show that *peb* overexpression results in ISC-to-EC differentiation. To resolve this contradiction, we focused on analyzing *peb* to better understand its function in the intestinal progenitors.

We knocked down (KD) or overexpressed (OE) *peb* in adult intestinal progenitors and used confocal microscopy to observe cellular defects. Intestinal progenitors were labeled by GFP expression, EEs were marked with prospero (*pros*) while ECs were recognized by their polyploid nuclei. Compared to control, the raw cell counts (including GFP-positive cells, EEs, and ECs) were lower in both the *peb-i* and *peb-OE*, with a decrease more prominently observed in the posterior midgut than the anterior region (**Figure 3E-G,I**). This similarity in the cell-count profile, because of opposite perturbations of the same gene, is surprising and could be a reason why previous studies arrived at different conclusions. We reasoned that the differences between the opposite perturbations might be clarified if we refined our parsing of cell types. Although mature ECs are polyploid and can be readily distinguished from other cell types, premature ECs undergoing endocycling can be hard to discern from ISCs. Additionally, ECs that have rapidly differentiated from ISCs can be GFP-positive as a result of the perdurance of the protein expressed in the progenitors. Thus, we measured nuclei area as a non-biased way to profile ISC/EB differentiation. We observed a decrease in the nuclei size of GFP-positive cells in *peb-i* when compared to controls (**Figure 3H**). In contrast, *peb-OE* resulted in an increase in nuclei size of GFP-positive cells (**Figure 3J**). These results suggest that lowering *peb* levels suppressed ISC differentiation into EC whereas overexpression pushes ISC towards EC lineage. We also observed some differences between the nuclei size of anterior and posterior cells in the midgut (**Supplementary Note 2**). The accumulation of mature ECs in *peb-i* is consistent with a previous study which showed that blocking newborn ECs from forming increased the ploidy of existing ECs ^28^.

### Applying Haystack prioritization-validation on Drosophila larval blood

*Drosophila* hemolymph consists of three populations of blood cells or hemocytes: macrophage-like plasmatocytes (PM), platelet-like crystal cells (CC), and giant-cell like lamellocytes (LM), which express the known marker genes *NimC1*, *PPO1/2*, and *Atilla*, respectively ^29^. Recent advances in scRNA-seq methods have refined our understanding of the fly blood lineages, with the identification of additional marker genes, sub-populations, and various states within these three cell types ^6,30–32^. Using Monocle 3-based lineage analysis ^33^, the original study had determined that a putative oligopotent source subpopulation within PMs gives rise to the three main lineages ^6^. This is consistent with previous experimental evidence which suggests that both CC and LM populations can emerge from pre-existing plasmatocytes^34–36^. However, the lineage-determining TFs that promote the differentiation of PMs to other lineages, especially toward the LM lineage, are poorly characterized.

To identify TFs that are active along the *Drosophila* blood cell lineages, we applied Haystack on blood cell scRNA-seq pertaining to larvae upon wounding, which is sufficient to activate blood cells and induce LMs ^637^ (**Figure 4A**). For simplicity, PM clusters from the original study have been combined into two groups, PM^early^ and PM^late^, corresponding to the early (oligopotent) and late-stage (mature) PMs, respectively. The CC and LM cell-types are also subdivided into two clusters each based on the expression levels of their respective marker genes, *PPO2* and *Atilla* (**Figure S4A**). We separately analyzed the three lineage trajectories, each with its source set at oligopotent PMs (PM^early^, in blue) (**Figure 4B**). For the PM^early^ → CC lineage, Haystack efficiently prioritized four source TFs: *Dp*, *Jumu*, *Lz*, and *Myc* (**Figure 4C**). The Notch target genes *Lozenge* (*Lz*) and *Myc* are known regulators of CC development and lineage commitment ^38–41^. Also, recent evidence indicates that loss of *Jumu* decreases CC numbers ^37,42^. These data indicate that Haystack can efficiently prioritize TFs that are known to control terminal differentiation of blood cells. For the PM^early^ → LM lineage, Haystack prioritized two novel source TFs *Xbp1* and *CG3328* (**Figure 4C, S4B,E**), neither of which have been previously implicated for this lineage. We note that the human ortholog of *Xbp1* was also identified in the Haystack analysis of human leukemia data (**Figure 2F**), which suggests an evolutionarily conserved role for this TF in blood cell differentiation.

**Figure 4.**
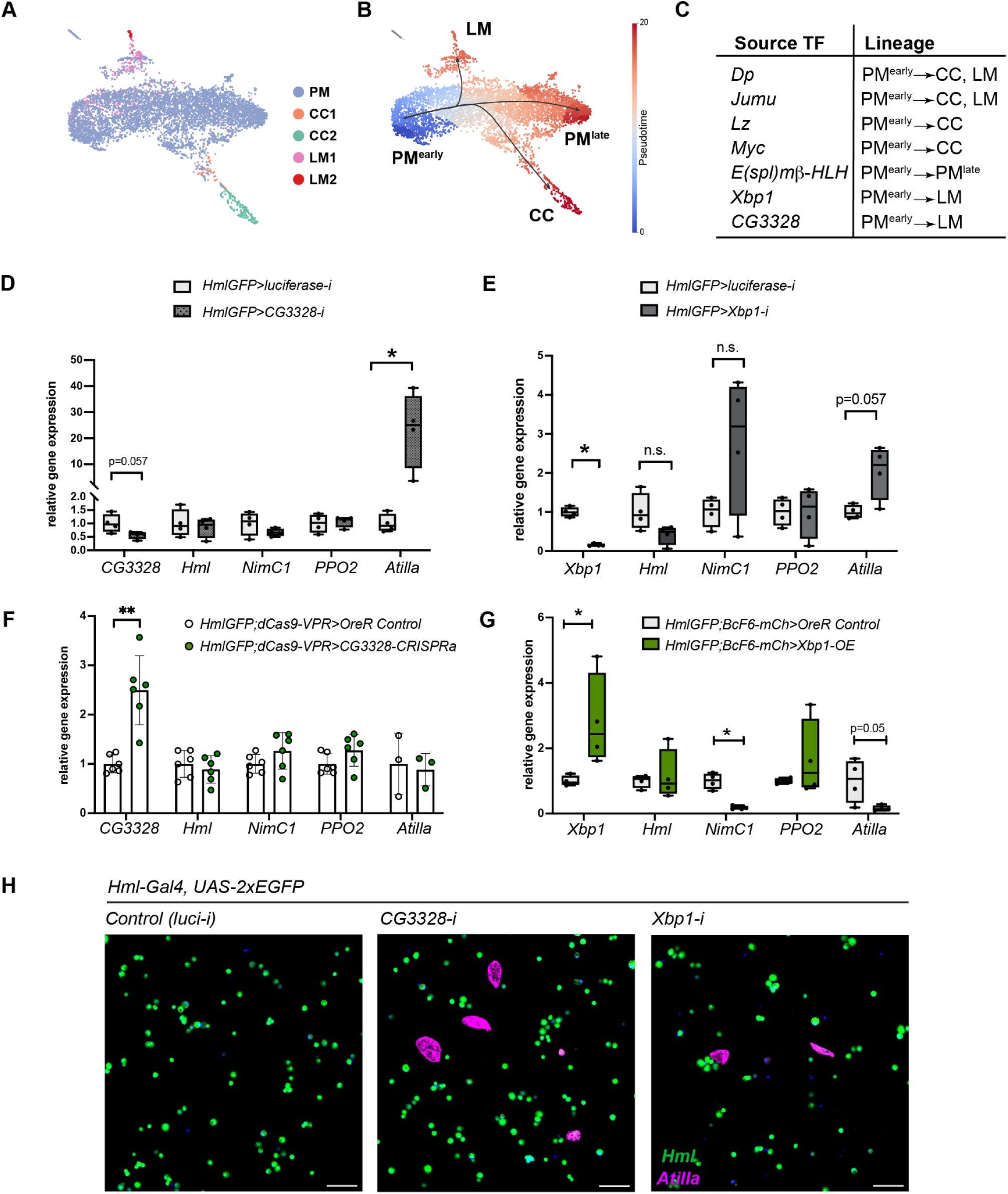
Haystack identifies novel TFs in the Lamellocyte lineage of Drosophila blood. **A.** *Drosophila* larval blood scRNA-seq^6^ shows plasmatocyte (PM), crystal cell (CC), and lamellocyte (LM) clusters. CC1 and LM1 represent putative immature while CC2 and LM2 represent mature CCs and LMs, respectively. **B.** Monocle 3-based pseudotime analysis shows that three lineages (PM^late^, CC, and LM) emerge from the oligopotent PM^early^ cluster. **C.** Table representing source TFs identified by Haystack. **D-E.** qRT-PCR analysis of larval blood upon knockdown of *CG3328* (D) and *Xbp1* (E) shows an increase in the LM marker gene *Atilla*. Non-parametric multiple t tests were used and n.s. and * represent not significant and *P*<0.05, respectively. N=4 biological replicates. **F-G.** qRT-PCR analysis of larval blood upon overexpression of CG3328 (F) and Xbp1 (G) shows respectively no change or a decrease in the LM marker gene Atilla. Non-parametric multiple t tests were used and n.s. and * represent not significant and P<0.05, respectively. N=4 biological replicates. **H.** Confocal imaging of blood cells derived from *HmlGFP>luci-i*, *CG3328-i*, and *Xbp1-i* larvae. Scale bar = 50um.

We focused on the LM lineage, as our understanding of the regulators of LM differentiation is limited. Hence, we knocked-down or overexpressed *CG3328* and *Xbp1* (**Figure 4D-G**) in blood cells using the *HmlD-Gal4, UAS-2xEGFP* driver (referred to as *HmlGFP* henceforth), where EGFP marks most PMs ^43^. To identify changes in blood cell lineages upon perturbation of *CG3328* and *Xbp1*, we first performed qRT-PCR on total RNA derived from circulating plus sessile blood cells of the various genotypes. RNAi-mediated knockdown of *CG3328* resulted in over 20-fold induction of the LM marker gene *Atilla*, while the expression of *NimC1* tended to be downregulated (**Figure 4D**), indicating a shift from PM to LM lineage. However, the changes in *PPO2* expression were commensurate with changes in *Hml* expression (**Figure 4D**), suggesting that CC differentiation is not impacted by *CG3328*. To address the role of gain-of-function of *CG3328*, we utilized CRISPR-mediated activation (*CRISPR^a^*) approach using the *HmlD-Gal4, UAS-EGFP; UAS-dCas9* (*HmlGFP;dCas9*) driver. However, increasing the levels of *CG3328* in PMs did not affect the expression of any of the lineage marker genes (**Figure 4F**). With regards to the perturbation of *Xbp1*, we observed a subtle increase in *Atilla* expression (**Figure 4E**). Interestingly, the expression levels of *NimC1*, a marker for mature PMs, tended to be upregulated suggesting that *Xbp1* regulates both PM^late^ and LM lineages. On the other hand, overexpression of *Xbp1* resulted in decreased expression patterns of *NimC1* and *Atilla*, while *Hml* and *PPO2* remain unchanged compared to *OreR* controls (**Figure 4G**). To further validate our findings from the qRT-PCR data pertaining to the role of these two TFs in regulating the LM lineage, we performed confocal imaging of blood cells in all the genotypes. As expected, the RNAi-mediated knockdown of *CG3328* resulted in an increase in LMs (**Figure 4H, S4C,D**) and knockdown of *Xbp1* showed a subtle increase in LMs (**Figure 4H, S4F,G**) compared to *luciferase-i* controls. Of note, our confocal imaging data corroborate the qRT-PCR data pertaining to the LM lineage. Lastly, besides the LM lineage, we also tested the role of *E(spl)mbeta* in the PM^early^ → PM^late^ lineage, as predicted by Haystack (**Figure S4H**). qRT-PCR analysis shows that overexpression of this TF decreased the expression levels of *NimC1* (**Figure S4I**), while the cell type compositions (of PMs and CCs) remain unchanged with no detection of Atilla+ LMs (**Figure S4J**). Altogether, these results indicate the predictive power of Haystack in shortlisting biologically relevant TFs for downstream lineage analyses.

Put together, Haystack helped accomplish efficient discovery by prioritizing-validating multiple causal TFs across two distinct tissue types. Leveraging it also allowed us to resolve a previously contradictory biology of *peb* in *Drosophila* gut lineage, in addition to the identification of two TFs, *CG3328* and *Xbp1*, as playing a causal role in regulating the LM lineage in *Drosophila* blood.

### Haystack prioritize-validates source-target TF pairings

Towards identifying signaling cascades of TFs, we next used Haystack to identify TFs localized to the end-points of trajectories (i.e., target TFs) in *Drosophila* gut differentiation. Using the cisTarget database, we limited our analysis of TFs that were putative transcriptional targets of the 8 source TFs described previously. We identified a total of 54 that satisfied this criterion and further narrowed them down to 13 TFs that localized to cells at the three endpoints. Although not all target TFs were downstream of each of the 8 source TFs, we measured the levels of all 13 target TFs under the 12 “source” perturbations (7 overexpression; 5 knockdown) to assess the validity of our putative predictions; we also measured all 8 source genes to confirm the perturbations. This amounted to a total of 252 observations (**Figure S3**). Among the 24 perturbations tested for putative source-target pairings, 14 (58%) were correct. That is, down regulating a source TF led to a decrease in a putative target TF or vice versa for overexpression. One of the novel source-target pairs was peb>*Myc*. *Peb-i* resulted in a decrease in *Myc* levels, whilst *peb-OE* increased the levels of *Myc*. This is consistent with previous studies of Myc function in the midgut where the overexpression of *Myc* increases nuclei size and knockdown of *Myc* reduces ISC numbers^44,45^. These phenotypes are remarkably similar to what is observed in *peb* perturbations. Our results indicate that our prioritization algorithm can accurately identify TF-TF source-target pairs (**Figure S5**).

## Discussion

Single-cell genomics, accompanied by computational analyses that generate a multitude of hypotheses^46^, has led to an explosive growth of data. However, more data does not imply more insight— our knowledge of the causal mechanisms of cell development has not kept pace with such data explosion. Currently, a key bottleneck in research progress is validating causal hypotheses generated from observational scRNA-seq studies. In this study, we developed a hybrid computational-biological method, Haystack, to prioritize crucial transcription factors (TFs) involved in differentiation for downstream validation. Employing Haystack on scRNA-seq datasets derived from mouse gut and human leukemia revealed that 75% and 84% of predictions, respectively, had literature supporting their function. Haystack’s high hit rate was also observed with the *Drosophila* datasets. In the fly gut, all of the 8 source TFs predicted to be involved in ISC differentiation (hit rate of 100%), and 5 of the 7 source TFs prioritized in blood cell differentiation (hit rate of 71%), had measurable qRT-PCR changes following genetic perturbation; imaging-based validation confirmed all source TFs shortlisted after qRT-PCR. This is a substantial improvement when compared to previous studies where a RNAi screen of essential genes in *Drosophila* ISCs achieved a hit rate of 6.4%^25^ and misexpression screens for defects related to blood cell composition yielded hit rates of ~30.7% and ~34.3%^47,48^. Our results show that Haystack can effectively identify and prioritize TFs for in vivo validations.

### Effective hypothesis prioritization drastically reduces time and money costs

Effectively prioritizing TFs for validation is crucial for understanding cell differentiation and lineage determination. Most scRNA-seq studies use algorithms that predict lineage trajectories and reveal an abundance of hypotheses that often exceed what can be tested in vivo. The cost associated with validation takes the form of time and money that is spent on validating TFs implicated in differentiation. In our mouse gut analysis, prior to Haystack prioritization we identified 98 TF regulons as implicated in cell differentiation. To validate the in vivo function of all these TFs in mouse studies, we estimate that a lab may spend tens of thousands of dollars and hundreds of person-hours. For model organisms like *Drosophila*, where one generation of breeding takes ~10 days, the time saved outweighs monetary benefits. The rate-limiting step in *Drosophila* validations is the phenotypic analysis: preparation of tissue, acquisition of confocal images, and image analysis are all laborious procedures (~5-10 hours for each assay). The high hit rate of Haystack’s prioritization algorithm drastically reduces validation efforts that are required to achieve the same nominal rate of causal conclusions.

### The novel use of optimal transport to rank the cost function of TF activity distributions

Rather than de novo GRN inference, the key algorithmic contribution of Haystack is effectively prioritizing (i.e. re-ranking) existing GRN predictions by combining multiple perspectives. While there are several computational methods that calculate TF module activity in clusters and others that infer pseudotime trajectory, none, to our knowledge, have attempted to combine the two categories that we employed in Haystack. Here, we use SCENIC to estimate TF module activity, but other approaches (e.g., SCENIC+ or CellOracle) could also be used. Similarly, RNA velocity based approaches (rather than pseudotime) can be used to compute the differentiation trajectory.

A key advantage of the non-parametric nature of OT is its robustness to non-uniform distributions of pseudotime values. Since pseudotime is a score that emerges from a graph-theoretic analysis of the single-cell landscape, there is no guarantee that cells will be uniformly distributed across it. Compared to an approach that would identify early TFs by simply correlating with pseudotime scores, our OT based formulation is more robust to non-uniform distributions. Previously, Schiebinger et al. have used OT to reconstruct cell lineages along a differentiation trajectory to infer ancestor-descendent cell fates^49^. Our work demonstrates how OT can also be leveraged for experiment design in single-cell genomics and used to guide an active learning approach.

### Haystack captures per-cell TF activity independent of cell type discretization

Haystack is applicable to a single scRNA-seq study and does not require cell type annotations, so it can be used in cases where unknown or transient cell types are involved. A limitation of many existing methods is a need to specify clear stages of differentiation, either as multi-time course study, or by annotating discrete cell types along the differentiation process. This imposes substantial data needs– in a recent study, Qiu et al. sought to identify TFs that specify cell types that emerge during mouse gastrulation by integrating ~480 scRNA-seq datasets (comprising ~1.6 million cells) over 19 stages spanning from E3.5 to E13.5^50^. In contrast, the use of OT metrics allows Haystack to obviate the need for cell type discretization and annotation by making inferences at the level of individual cells, rather than cell types.

### Concluding Remarks

The active learning and OT framework of Haystack represents a general approach to prioritize and validate lineage-determining regulatory factors. For instance, the framework can be extended to other forms of aggregate data such as gene modules affected by metabolic pathways or microRNAs. The distribution of these modules, as a function of pseudotime, can then be compared with a baseline and ranked with OT cost functions. These rankings will generate regulatory hypotheses that can undergo the same iterative testing as shown in our study. In all, these potential experiments and our results underscore the importance of using principled mathematical approaches and active learning frameworks to identify causal biological conclusions from big data.

## STAR Methods

### Trajectory analysis and transcription factor (TF) module identification

Trajectory analysis and pseudotime computation of the scRNA-seq data was used to estimate the differentiation time-course; in the case of multiple branching trajectories, we repeat our analysis for each branch separately and aggregate the results. Haystack can be applied with a preferred pseudotime (or RNA velocity) estimation program; here we present results with SlingShot^21^, diffusion pseudotime^51^, and Monocle 3^33^, choosing the method used in the original scRNA-seq study whenever available. The optimal transport analysis in Haystack does not require any knowledge of the cell types in the dataset and the differentiation stage of a cell is inferred solely from pseudotime analysis. When cell types are known, they may be used to limit Haystack analysis to a lineage of interest. TF regulons are mapped over the differentiation time-course, where regulons are inferred from the cisTarget database of TF binding sites.

### Characterizing TF localization through optimal transport

To identify TFs that are active in just one differentiation stage and not broadly active, we arrange cells along a one-dimensional pseudotime axis (**Fig. 1a**). Each TF activity is characterized as a probability distribution along the pseudotime axis, computed from the histogram of per-cell regulon activity indicators (for each regulon, SCENIC^2^ reports the cells with statistically significant activity of the regulon). We also compute the baseline probability distribution (i.e. histogram) of all cells along this axis. Using optimal transport (OT), we compute for each TF the distance between its distribution and the baseline. OT is a mathematical formulation for measuring the distance between two probability distributions under some cost function. In our context, the cost between two cells corresponds to the difference of their score along pseudotime axis, capturing the distance between their differentiation stages; other measures like RNA velocity could be similarly incorporated. More formally, let *p_0_(t)* be the probability distribution of cells along the pseudotemporal axis *t*. Here, we normalize *t* to range [0,1] and approximate *p_0_(t)* from the histogram of cell counts along pseudotime. Similarly, let *p_G_(t)* be the probability distribution corresponding to the activation of the regulon corresponding to TF *G*. We compute the Wasserstein distance *d_W_(p_0_,p_G_)* between the two distributions *p_0_* and *p_G_* by solving the transport problem:

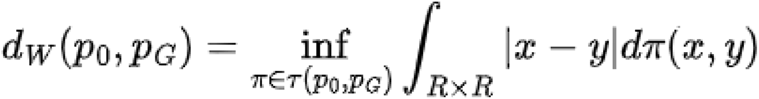

Where τ(*p*_o_ *p*_g_) is the set of probability distributions on *R* × *R* whose marginals are *p_o_* and *p_G_*, respectively. In this setting, the cost function *c(x,y) = |x-y|* is simply the pseudotime distance between two points. We computed the Wasserstein distance using the Python package *scipy* (version 1.6.0)

Intuitively, TFs that are active only in a limited region of the time-course will be at a substantial OT distance from the baseline. We note that the OT metric captures not only the differences of mean between distributions (first moment) but also differences in higher moments: e.g. even if the probability distribution of a TF has the same mean as the baseline, if the former is concentrated around this location (i.e. has a lower variance than the baseline), the OT distance can be large. The use of OT offers key advantages over alternative approaches. In OT, the empirical probability distribution of TFs over cells is directly computed upon, so we do not need to assume that the underlying distribution obeys specific properties (e.g. those required by statistical tests like the t-test). Unlike some metrics over probability distributions (e.g. mutual information or Jensen-Shannon distance)^52^, the OT formulation incorporates the concept of cost which allows us to account for the pseudotemporal distance between two TFs (or between a TF and the background).

If a TF has high activity in two separate pseudotime regions distinct from the baseline (e.g. at both the start and end of the time-course), the OT metric may also score such non-localized bimodal distributions highly. To address such potential false positives, we incorporate an additional measure of TF localization: we represent each cell as feature-vector of TF activations and identify the features (i.e. TFs) that are most informative of the cell’s pseudotime score. We built upon Schema^53^, a metric learning approach for feature selection in multimodal single-cell data, to solve a quadratic program to identify TFs whose activation is most informative of the cell’s pseudotime score. We ensembled the two approaches by converting their outputs to rank-scores of TFs and computing a weighted combination of the two. Small-scale explorations indicated that results were robust to the choice of weight around 0.5, which we have chosen for all results presented here. We then limited our analysis to the top-third of the TFs.

### Identifying lineage-determining TFs

Lineage-determining TFs typically have more than one downstream target^9^, with their ability to influence a broad transcriptional program being key to fate determination. Accordingly, we queried the cisTarget database^54^ to identify TF pairs (TF_source_–TF_target_) where the binding site of TF_source_ was found upstream of TF_target_, suggesting that TF_source_ might regulate TF_target_. From the shortlist of well-localized TFs, we considered every pairwise combination of TFs (say, TF_1_ and TF_2_) such that TF_1_ is active earlier in the pseudotime landscape than TF_2_ and with the binding site of TF_1_ upstream of TF_2_. We then applied OT to compute the pseudotime distance between the two TFs. From these pairs, we extracted the subset corresponding to high OT distance. The source TFs in these pairs are the candidate TFs we generate as the initial set of high-confidence hypotheses prioritized for experimental validation. Within this set, we rank TFs by the number of well-separated pairs in which the TF occurs as a source.

### Estimating cell-type composition from qRT-PCR assays of marker genes

Quantitative Real Time Polymerase Chain Reaction (qRT-PCR)-based validation provides a medium-throughput validation that is more efficient than imaging and phenotypic studies. For each of the shortlisted TFs, we perform perturbation experiments (e.g., for *Drosophila* gut and blood studies, we overexpressed or knocked down the TFs of interest). From the original scRNA-seq study, we identified the cell types/clusters of interest and applied differential expression analysis to identify a limited set of markers (typically, 1-3) per cell type. In our *Drosophila* assays, this resulted in less than 15 markers in total, and thus amenable to a single qRT-PCR study. We assayed these markers to assess changes in cell type composition as a result of the perturbation, by comparing qRT-PCR values in the wild type tissue against the overexpressed/knocked-down tissue. Furthermore, since the markers are the same for each perturbation, the same qRT-PCR primers and setup can be used across all perturbations in parallel, speeding up the process.

A confounding issue in estimating cell-type composition changes using qRT-PCR is that a perturbation may vary the overall proliferation rate of the tissue vis-a-vis the organismal background. Since qRT-PCR cycle threshold (CT) values are typically normalized against the background CT value of a housekeeping or ribosomal gene, this can confound cell type composition analysis. Accordingly, we introduce a fold-change metric to adjust for this confounding factor and robustly recover estimates of cell type compositions: we first compute robust estimates of cell type expression by averaging the markers for each cell type. We then choose one cell type’s abundance as the baseline (typically, the progenitor cell type), and express all other cell type abundances as a ratio against this baseline. Wild-type and perturbation lines can now be compared using a fold-change metric on this ratio to identify which, if any, differentiated cell types have increased or decreased.

### Applying Haystack on mouse gut and acute myeloid leukemia cells

We applied Haystack on the scRNA-seq studies of mouse gut differentiation^10^ (GSE152325) and human acute myeloid leukemia (AML) cells^11^ (<u>10.5281/zenodo.3345981</u>). We used these datasets to first reconstruct cell lineages (diffusion pseudotime algorithm for the mouse gut as in the original study; Monocle 3 for the AML study) and identified TF modules along the trajectories. In the mouse gut, intestinal stem cells (ISC) differentiate along four lineages: enteroendocrine (EE) cells, tuft cells, goblet cells and enterocytes (ECs). Applying Haystack on each of the lineages, we identified the source TFs localized to or near the ISC cluster. Interestingly, when we visualized individual TFs using transcript levels alone (eg. *Ezh2*, *Sox4*, *Sp5* and *Nr3c1*) (**Figure 2C**), a diffuse expression pattern was observed. Instead, visualizing TF modules revealed that the same diffusely expressed TFs had very localized activity. We aggregated these results across the lineages by counting the total number of downstream targets per source TF and ranked them within each lineage. Of the 24 putative source TFs predicted by Haystack, 75% had substantial support in the literature for their involvement in the development of the gut, most of which were related to differentiation (see **Table S1**).

For the human AML dataset, the pseudotime analysis revealed that hematopoietic stem cells (HSC) differentiate into dendritic cells, erythrocytes, NK T cells and monocytes in a linear trajectory. We found that TF modules exhibited clearly localized activity while the raw gene expression was diffusely expressed (e.g., *Etv6*, *Xbp1*, *Irf1* and *Runx3*) (**Fig. 1g**). We identified a total of 25 high-confidence TF predictions, ~85% of which had substantial evidence in the literature to support involvement in blood function (see **Table S2**).

### Systematic benchmarking of Haystack in the BEELINE framework

The BEELINE framework is a set of curated synthetic and experimental datasets (with corresponding ground-truth annotations and benchmarking software) for systematically evaluating GRN inference methods under uniform conditions^5^. Among the 12 GRN inference methods that were originally evaluated, PIDC^16^, SCODE^15^, Genie3^55^ and GRNBoost2^56^ were the top performers, especially on experimental data (Figure 6 of the original study). Here we chose PIDC and SCODE as benchmarks and excluded Genie3 and GRNBoost2 since they are indirectly used by Haystack as subroutines. Haystack leverages SCENIC to robustly infer TF activation and, in turn, SCENIC uses Genie3 or GRNBoost2 (depending on the user’s preference) to establish preliminary TF–target edges before refining them. Additionally, we also included SINGE^17^, which performed well on synthetic BEELINE data. We did not evaluate SCENIC separately here since it does not incorporate the differentiation trajectory or explicitly rank TFs, making it infeasible to evaluate the precision of its top hits.

We focused our evaluation on experimental scRNA-seq datasets since our goal is to enable efficient wet-lab perturbations. Synthetic datasets are not reflective of real-world utility due to their small gene sets, simple dynamics, and low noise. From the BEELINE framework, we obtained scRNA-seq data from five differentiating tissues: 1) mouse hematopoietic stem and progenitor cells (mHSC), 2) mouse embryonic stem cells (mESC), 3) mouse dendritic cells (mDC), 4) human mature hepatocytes (hHep), and 5) human embryonic stem cells (hESC). The mHSC dataset consisted of trajectories with multiple branches and was further subdivided into three subgroups: mHSC-E (erythroid), mHSC-L (lymphoid) and mHSC-GM (granulocyte-monocyte).

Existing GRN inference methods build a regulatory graph, encompassing both TFs and genes, directly from the provided input of expression profiles. Following the BEELINE framework, we applied each method on expression data of all TFs and the 1,000 most variable genes in each tissue. Where required, pseudotime information was also provided as an input. From the ranked list of gene–gene edges reported by each method, we extracted all edges that originate from a TF. Next, TFs were ranked based on their score computed as the sum of unsigned edge-weights (i.e., both activation and repression) originating from itself. This approach is similar to the TF ranking procedure described in Matsumoto et al.^15^ The precision of top 10, 20 or 30 TFs by this ranking scheme were evaluated against ground truth annotations (**Figure 2G, S1A-B**). We also tested an approach where each method was applied on just the set of TFs so that all reported GRN edges would be TF–TF; this approach performed substantially worse and was abandoned.

For each scRNA-seq dataset, the BEELINE framework also offers ground-truth regulatory associations based on cell-type specific and non-specific ChIP-seq data. We converted both types of ChIP-seq data separately into a TF ranking as above, selecting the top 100 TFs as the gold standard to compare predictions against. Unlike the original study, we did not use the STRING database of protein associations^57^. Since the proteomic data in STRING contains an uneven collection of predicted (e.g., coexpression), indirect (e.g., homology-based) and direct physical interaction edges, we believe it is not an appropriate ground-truth benchmark for evaluating transcriptional regulation.

In our evaluations, we report Haystack results derived from a ranked list of source-only TFs as well as rankings that combine both source and target TFs (**Figure S5**). While we expect source TFs to be the focus of perturbational assays, we also report the combined source-and-target rankings to ensure a fair comparison with the baseline methods, which may not make distinctions between a source versus a target TF. We also explored Haystack’s performance at different choices of a key hyperparameter by varying the cutoff threshold for selecting well-localized TFs. Haystack’s outperformance was robust to this choice (**Figure S1c-f**).

### Benchmarking Haystack on mouse gut and AML datasets

#### Generating a reference gene set by text mining

A standard approach to evaluating gene prioritization methods is to assess the subset of genes they prioritize against a reference list of genes documented to be involved in the process of interest. To our knowledge, for mouse gut or human AML differentiation no such reference gene sets are available. We therefore approximated the reference gene sets for the two tissues by text mining of research studies. Though these reference sets have not been derived by manual curation, we believe that our systematic approach results in unbiased errors, making these sets a useful validation benchmark. For each tissue, a collection of relevant studies was first acquired by querying the Pubmed database for publications whose title or abstract indicate that the study likely corresponds to the tissue of our interest. The queries used were:

- Mouse gut: (“mouse”[Title/Abstract]) AND (gut[Title/Abstract] OR intestine[Title/Abstract]) AND (development[Title/Abstract] OR differentiation[Title/Abstract])
- Human AML cells: (human[Title/abstract]) AND (hematopoiesis[Title/Abstract] or “blood differentiation”[Title/Abstract] or leukemia[Title/Abstract]) AND regulation[Title/Abstract]

An alternative querying approach would have been to limit ourselves to MeSH (Medical Subject Headings) gene and function annotations. However, we found the MEDLINE annotation of Pubmed articles by MeSH terms to miss important genes for our biological processes of interest. For example, the following query which uses only MeSH terms to search for publications of *Ezh2* activity in mouse intestine development returned zero hits at the time of submission. In contrast, our approach identified four relevant publications (e.g., see Ezh2-related reference in **Table S1**)

- ((“Ezh2 protein, mouse” [Supplementary Concept]) OR (“Polycomb Repressive Complex 2”[Mesh])) AND (“Intestines/growth and development”[Mesh])

From the titles and abstract of the queried list of publications, we identified gene names. The set of protein-coding gene names and their synonyms was sourced from the NCBI Gene database. We excluded gene synonyms that are also common English words. For instance, for the *WASP actin nucleation promoting factor* we included the synonyms *IMD2, SCNX, THC, THC1, WASP*, and *WASPA*; but we excluded the synonym *WAS*. The final reference set for each tissue was chosen as the genes listed in at least two publications (**Table S4**).

#### Evaluating early precision with a gene enrichment metric

We computed the top TFs prioritized by Haystack as well as other methods (**Fig. 1i**). We evaluated three gene network inference methods: SCODE^15^, PIDC^16^, and SINGE^17^. Due to SINGE’s long run-time (10 days for a dataset with 3,000 cells), we applied it to a random subset of 3,000 cells in human AML tissue and 2,000 cells in mouse gut tissue. For each method we selected the top 30 hits (fewer if the method reported less than 30 TF hits). We also shortlisted TFs by a differential expression analysis, performing the Wilcoxon rank-sum test to identify TFs differentially expressed between progenitor and other cell types as annotated in the original studies. The requirement of a pre-curated annotation might be infeasible for a poorly-studied tissue; therefore, neither Haystack nor the gene network inference methods require such pre-curated annotations. For each gene set, its enrichment in the reference set was evaluated, computed as the fold-change over the expected enrichment of an equal-sized random set of protein-coding genes.

As another control, we computed the fold-enrichment of the set of all TFs in the species, finding this set to be enriched in both tissues. We interpret this as supporting our text-mining approach to extracting relevant studies: it would have been surprising if published studies of differentiation in these tissues did not highlight TFs more than other genes.

#### Haystack’s improvement over SCENIC

We also attempted to examine the additional accuracy of Haystack over SCENIC. Notably, SCENIC does not consider the differentiation landscape and TFs are not ranked by their likelihood of being a source. Nonetheless, we evaluated the full set of SCENIC regulons (93 in the case of mouse gut; 219 for human AML cells), computing their enrichment against equal-sized sets of random protein-coding genes. In both the human AML and mouse gut tissues, the set of SCENIC regulons is substantially more enriched than random (fold-enrichment scores of 6.11 and 5.67, respectively), indicating that its module discovery process does enhance the TF signal in the data. However, SCENIC by itself is not sufficient— it outperformed single-gene differential expression in mouse gut (score of 2.34) but underperformed the latter in human AML cells (score of 5.84). In comparison, the set of TFs prioritized by Haystack (scores of 6.81 and 8.12 in mouse gut and human AML, respectively) not only displayed higher enrichment than either SCENIC or differential expression, but also higher than the gene network inference methods (**Figure 2H**).

### Runtime and memory usage requirements of Haystack

We assume here that pseudotime trajectories have been already computed by the researcher’s method of choice. Once TF modules and pseudotime have been computed, the optimal transport computation in Haystack runs under 5 minutes and requires less than 8 GB of RAM. The preprocessing step of computing TF modules via SCENIC (https://github.com/aertslab/pySCENIC) is more time intensive: on a 10,000 cell dataset, computing TF modules using the pySCENIC Docker instance (using the GRNBoost2 sub-module and with parallelization enabled) required 22 minutes on a 24-core Intel Xenon 3.5 GHz server with peak memory consumption under 30 GB. The run-time of SCENIC increases with the number of cells and we therefore recommend sketching approaches^58^ to downsample datasets with hundreds of thousands of cells. We also recommend using GRNBoost2, rather than Genie3, as the subroutine inside SCENIC since the former is substantially faster.

### *Drosophila* stocks and culture

Flies were reared in humidified incubators at 25°C on standard lab food composed of 15 g yeast, 8.6 g soy flour, 63 g corn flour, 5 g agar, 5 g malt, 74 ml corn syrup per liter with 12/12 hr dark/light cycles. For all Gal80^ts^ (temperature sensitive) experiments, crosses were reared at 18°C. After eclosion, flies were kept at 18°C for 3 days before shifting to 29°C (permissive temperature) for 10 days. For all blood experiments, fly larvae of respective genotypes were grown on the standard lab food until late third larval instar (LL3) at 25°C.

The following stocks were obtained from the Bloomington *Drosophila* Stock Center (BL), DGRC (NIG) and FlyORF: *UAS-Luc-i* (BL36303), *UAS-peb-i* (BL28735), *UAS-peb* (BL5358), *UAS-Psi-i* (BL31301), *UAS-Psi* (BL16371), *UAS-drm* (BL7072), *UAS-Tet-i* (BL62280), *UAS-Mondo-i* (BL27059), *UAS-Mondo* (BL20102), *UAS-cnc* (BL17502), *UAS-Tet-i* (BL62280), *UAS-tbp-i* (NIG 9874R-1), *UAS-FoxK* (F000615), *UAS-Mondo* (F001398), *UAS-Xbp1* (BL60730), *UAS-E(spl)mbeta-HLH* (BL26675), *UAS-CG3328-i* (BL55211) and *UAS-Xbp1-i* (BL36755). The following Gal4 lines used to perturb genes in guts and hemocytes, respectively, were: *esg-Gal4* and *w[1118];Hml-Gal4.Delta,UAS-2xEGFP* (BL30140), hereafter referred to as *HmlGFP*. *Oregon R* (*OreR*) control flies were obtained from the Perrimon Lab stock. The *BcF6-mCherry* (a crystal cell reporter)^59^ stock obtained from Dr. Tsuyoshi Tokusumi (Schulz Lab), was crossed with the *HmlGFP* line to obtain *HmlGFP;BcF6-mCh* stock. To drive the expression of *CG3328* in blood cells in a Cas9-based transcriptional activation (*CRISPR^a^*) manner^60^, the *Hml-Gal4,UAS-EGFP;dCas9-VPR* (*HmlGFP;dCas9-VPR*) was crossed to *CG3328-sgRNA* fly line (BL80297).

### RNA extraction and qRT-PCR

#### Drosophila midguts

7-10 midguts were dissected in 1xPBS and homogenized in 300uL of TRIzol (ThermoFischer, cat# 15596-026) using RNase-free pestles. RNA was extracted using Zymo Direct-zol RNA MicroPrep kit (cat# R2060) and subsequently DNase-treated using Turbo DNA free (cat# AM1907). 400-450ng of the resulting RNA was reverse transcribed using

Bio-Rad iScript Select cDNA synthesis kit (cat# 708896) and SyBr green (cat# 1708880) based qRT-PCR was performed to determine the levels of gene expression. qRT-PCR primers were designed using FlyPrimerBank^61^. The efficiency of primers was determined by running qRT-PCR on serial dilutions of pooled cDNA. Only primers in the range of 85% to 110% efficiency were selected for further use. See Table S3.

#### Drosophila hemocytes

RNA isolation of larval blood was performed as described previously with minor modifications^6^, where hemolymph from ~15-20 larvae (or ~50 for a better yield) are sufficient for RNA isolation per biological replicate. For a detailed protocol regarding hemocyte isolation, RNA/cDNA preparation and qRT-PCR set up, see https://en.bio-protocol.org/prep1155

### Immunostaining and imaging

#### Drosophila guts

Whole midguts were dissected in PBS and fixed in 4% PFA in PBS at room temperature for 30 minutes. Fixed guts were washed once in 0.1% Triton X-100 in PBS (PBST), then blocked with a blocking buffer (0.1% Triton, 5%NDS in PBS) for 30 minutes at RT. Primary antibodies were incubated overnight at 4°C in the blocking buffer. Guts were washed 3x in the blocking buffer and incubated with secondary antibodies overnight at 4°C along with DAPI (1:1000 of 1mg/ml stock). After antibody staining, guts were washed 3 times in PBST and mounted in Vectashield antifade mounting medium (Vector Laboratories cat# H-1200). Tape was used as a spacer to prevent coverslips from crushing the guts. Antibody dilutions used were as follows: chicken anti-GFP (1:2000, Abcam cat# ab13970), donkey anti-rabbit 565 (1:2000, Molecular Probes cat# A31572), goat anti-mouse 633 (1:2000, Thermo Scientific cat# A-21240) and goat anti-chicken 488 (1:2000, Thermo Fisher Scientific cat# A-11039). Guts were imaged on a spinning-disk confocal system, consisting of a Nikon Ti2 inverted microscope equipped with a W1 spinning disk head (50um pinhole, Yokogawa Corporation of America) and a Zyla 4.2 Plus sCMOS monochrome camera (Andor).

#### Drosophila hemocytes

20 late third instar larvae (LL3) from each genotype were vortexed, and bled in 300 ul of Schneider’s media in a 9-well spot glass plate. Next, the media with hemocytes was transferred to 8-well chambered cover glass slide (VWR, cat# 62407-296) and the cells were allowed to settle at room temperature for ~30 min. Alternatively, 96-well glass bottom plates (Cellvis, cat# P96-1.5H-N) were also used to plate hemocytes from 10 LL3 larvae per biological replicate per well. Next, 4% (final concentration) paraformaldehyde (Electron Microscopy Services, cat# 15710) was added to each well with Schneider’s media and hemocytes and incubated on a rocker for 20 min. Later, the fixed hemocytes were washed three times with 1x PBS (Gibco, cat# 10010-023) and blocked with a blocking buffer (5% BSA in 1x PBS containing 0.1% Triton-X) for 10 min. The cells were incubated with 1:100 dilution of anti-Atilla L1abc antibody^62^ overnight at 4°C. The next day, the cells were washed three times with 1x PBS and incubated with corresponding secondary antibody (1:500 dilution, anti-mouse alexa fluor 633) for 1 h at room temperature. Finally, the cells were washed three times with 1x PBS and Vectashield containing DAPI (Vector Laboratories Inc., cat# H-1200) was added before imaging the cells using Nikon Ti2 Spinning Disk Confocal Microscope. Cell counts were performed on 3-4 independent regions of interest (ROIs) per well (biological replicate) captured by the Nikon Ti2 Spinning Disk Confocal Microscope or GE IN Cell Analyzer 6000 Cell Imaging System. All images were analyzed by Fiji ImageJ software^63^.

## Code and Data Availability

Python code and instructions for use of the Haystack framework are available at: https://cb.csail.mit.edu/cb/haystack/

Source data from the imaging and qRT-PCR assays is available upon request. The following public datasets were used in the analysis:

| Description | Reference | Accession/URL |
| --- | --- | --- |
| Human leukemia scRNA-seq | Petti et al. | [10.5281/zenodo.3345981](https://doi.org/10.5281/zenodo.3345981) |
| Drosophila midgut | Hung et al. | GEO: GSE120537 |
| Mouse gut differentiation | Bottcher et al. | GEO: GSE152325 |
| Drosophila blood | Tattikota et al. | GEO: GSE146596 |
| Beeline Benchmark Data | Pratapa et al. | https://doi.org/10.5281/zenodo.3378975. It includes scRNA-seq data from GEO datasets, GSE81252 (hHEP), GSE75748 (hESC), GSE98664 (mESC), GSE48968 (mDC) and GSE81682 (mHSC) |
| cisTarget | Aibar et al. | https://resources.aertslab.org/cistarget/ |

**Software**: Python packages scanpy (v1.4.6), scipy (v1.6.0), scikit-learn (v0.24.1), and schema_learn (v0.1.5.3) were used. The R packages Monocle (v3), SlingShot (v3.15) were used. SCENIC (v1.1.1-7) was also used. In addition, the Github repository of Beeline (v1.0, https://github.com/Murali-group/Beeline) was used.

## Acknowledgments

We thank the assistance provided by the Microscopy Resources on the North Quad (MicRoN) core and the GE IN Cell Analyzer 6000 Cell Imaging facility of the *Drosophila* RNAi Screening Center (DRSC) at Harvard Medical School. We thank the members of the Berger and Perrimon labs for helpful discussion and feedback. RS and BB were partially supported by NIH NIGMS R35GM141861. JSSL was supported by the Croucher fellowship for Postdoctoral Research from the Croucher Foundation. This work was supported by NIH NIGMS P41 GM132087 and vBBSRC-NSF (NP). NP is an investigator of Howard Hughes Medical Institute. This article is subject to HHMI’s Open Access to Publication policy. HHMI lab heads have previously granted a nonexclusive CC BY 4.0 license to the public and a sublicensable license to HHMI in their research articles. Pursuant to those licenses, the author-accepted manuscript of this article can be made freely available under a CC

## Supplementary Information

### Supplementary Note 1

#### Cost of perturbations

Consider a relatively simple scenario, where one needs to analyze 10 homozygous mice that can only be generated from breeding heterozygous (het) animals. A total number of 40 offspring would be required to obtain these 10 (25%) homozygotes. Assuming a breeding female averages 6 pups/litter, a total of 7 het breeding pairs (7 females and 7 males; 7*6=42) would be required to generate one experimental cohort in one round of breeding (3 months). If het animals are not available, rearing wild type animals with a het animal generates ~50% het offspring. Thus, to generate 14 het mice would require 28 offspring. As such, ~5 WT x het breeding pairs are needed. Altogether, starting with 5 WT X het breeder pairs, the 10 homozygous mice would be ready in about 6 months. For just one experiment, approximately 80 mice will be generated. It would cost ~$2800 for housing the mice with the assumption of $1.25 per diem per animal (Boston University, https://tinyurl.com/yc5akwu8). This is an underestimation since in reality there are costs associated with reagents for genotyping and extra crosses that buffer for unsuccessful breedings. Additionally using qRT-PCR as a proxy for cell-type composition bypasses the need to buy costly reagents like antibodies or *in situ* probes to label cell-types. This highlights why selectively prioritizing high-confidence TFs for perturbation experiments is crucial for reducing cost.

### Supplementary Note 2

#### Applying Haystack on *Drosophila* midgut

The fly midgut consists of a monolayer of absorptive enterocytes (ECs) and secretory enteroendocrine cells (EEs) that are replenished by self-renewable intestinal stem cells (ISCs)^19,20^. Previously, we had performed scRNA-seq on fly whole guts and identified a total of 22 clusters mainly consisting of finer sub-classifications of EEs and ECs^7,18^. Some of these subtypes are distinguished by their spatial location whereas others were intermediate states between ISC/EB and a specific terminal state^18^. We applied Haystack on this scRNA-seq dataset. To avoid the confounding effects of unknown cell clusters, we limited our analysis to the known midgut cell-types; including EEs, ISC/EBs, <u>a</u>nterior ECs, <u>d</u>ifferentiating ECs, <u>m</u>iddle ECs and <u>p</u>osterior ECs (**Fig. S2a**). Using SlingShot^21^, we mapped individual cells onto a pseudotime trajectory consisting of three lineages with the starting point set as ISC/EB and end points as EE, aEC or pEC (**Fig. 2a,b**). By mapping TF module activity along these trajectories, Haystack identified eight TFs that were localized to the ISC/EB starting point (Source TFs). These included *Forkhead box K (FoxK)*, *cap-n-collar (cnc)*, *pebbled (peb)*, *P-element somatic inhibitor (Psi)*, *drumstick (drm)*, *TATA binding protein (Tbp)*, *Ten-Eleven Translocation family protein (Tet)* and *Mondo* (**Fig. S2b**).

As a broadly applicable intermediate validation, we measured mRNA levels (by qRT-PCR) of gene markers of specific cell types to approximate the cell-type composition. Using differential gene analysis, we selected a total of 11 markers that were distinctly expressed in ISC/EB (*Dtg, N, LanB1*), aEC (*CG6295, Npc2f*), pEC (*LManVI, Gs2, Mur29B*) or EE (*esg*, *AstC, IA-2*) (**Fig.S2b**). In the original Hung et al. study, these markers had been identified for individual cell clusters using a differential expression analysis in Seurat^27^ and for each cell type, we chose a combination of markers such that all sub-clusters of a cell type were covered. We knocked-down or overexpressed the eight TFs specifically in adult ISC/EB with available reagents. Among these, we found five RNAi lines (*peb-i, Psi-i, Tet-i, Mondo-i, cnc-i*) and seven overexpression (OE) lines (*peb-OE, Psi-OE, FoxK-OE, drm-OE, two Mondo-OE, cnc-OE*) that successfully reduced or increased, respectively, the gene of perturbation when assaying mRNA extracted from the whole midgut (**Fig. S2d**). We measured the levels of the 11 markers for each perturbation, which amounted to a total of 132 observations. We calculated the fold change (FC_RpL32_) in comparison to a control, using *RpL32* as a reference. In all cases, perturbation led to at least one, and up to 11, significant fold change(s) in marker gene expression. The FC_RpL32_ in markers gave us a proxy for the cell-type composition of the gut. For example, knockdown of *Psi* in intestinal progenitors caused a decrease in EE and pEC markers whilst ISC/EB and aEC markers remained unchanged (except for *Npc2f*). This suggests that *Psi* is involved in promoting the differentiation of progenitors to EEs and ECs in the posterior midgut (Fig. 2d). *Drm* overexpression caused an increase in ISC/EB markers with no changes in most terminal cell-type markers other than an increase in *Mur29B*. This suggests that *drm* promotes ISC proliferation and may not be involved in differentiation (**Fig. S2d**).

To overcome the caveat that terminal markers are confounded by the proliferation of progenitors, we used the EC–ISC mean marker ratio to compare our perturbations with the control. For *peb* perturbations, we knocked-down or overexpressed *peb* in adult intestinal progenitors and used confocal microscopy to observe cellular defects (see **Fig. 2e,f**). Intestinal progenitors were labeled by GFP expression and ECs were recognized by their polyploid nuclei (**Fig. S2e-f**). Compared to control, the raw cell counts (including GFP-positive cells, EEs, and ECs) were lower in both the *peb-i* and *peb-OE*, with a decrease more prominently observed in the posterior midgut than the anterior region (**Fig. S2e-f**). This similarity in the cell-count profile, because of opposite perturbations of the same gene, is surprising and could be a reason why previous studies^25,26^ arrived at different conclusions. We reasoned that the differences between the opposite perturbations might be clarified if we refined our parsing of cell types. Although mature ECs are polyploid and can be readily distinguished from other cell types, premature ECs undergoing endocycling can be hard to discern from ISCs. Additionally, ECs that have rapidly differentiated from ISCs can be GFP-positive as a result of the perdurance of the protein expressed in the progenitors. Thus, we measured nuclei area as a non-biased way to profile ISC/EB differentiation. In the anterior region, *peb-i* or *peb-OE* in intestinal progenitors resulted in no change in the mean nuclei area. The same perturbations, in the posterior midgut, caused a statistically significant but moderate increase in the mean nuclei area. Interestingly, differences could be observed by looking at the frequency distribution of the nuclei area. When compared to control, *peb* knockdown displayed a higher percentage of small nuclei cells (progenitors), a reduction in endocycling ECs, and an increase in large ECs in the anterior midgut. In the same region, *peb-OE* showed a reduction in the percentage of progenitors. In the posterior midgut, *peb-i* caused a decrease in early endocycling ECs but an increase in the late and mature ECs. *Peb-OE* causes a notable decrease in the proportion of progenitors and an increase in large ECs.

#### Using Haystack to predict source-target TF pairings

Towards identifying signaling cascades of TFs, we next used Haystack to identify TFs localized to the end-points of trajectories (i.e., target TFs) in *Drosophila* gut differentiation. Using the cisTarget database, we limited our analysis of TFs that were putative transcriptional targets of the 8 source TFs described previously. We identified a total of 54 that satisfied this criterion and further narrowed them down to 13 TFs that localized to cells at the three endpoints. Although not all target TFs were downstream of each of the 8 source TFs, we measured the levels of all 13 target TFs under the 12 “source” perturbations (7 overexpression; 5 knockdown) to assess the validity of our putative predictions; we also measured all 8 source genes to confirm the perturbations. This amounted to a total of 252 observations. Among the 24 perturbations tested for putative source-target pairings, 14 (58%) were correct (**Fig. S3**). That is, down regulating a source TF led to a decrease in a putative target TF or vice versa for overexpression. One of the novel source-target pairs was peb>*Myc*. *Peb-i* resulted in a decrease in *Myc* levels, whilst *peb-OE* increased the levels of *Myc*. This is consistent with previous studies of Myc function in the midgut where the overexpression of *Myc* increases nuclei size and knockdown of *Myc* reduces ISC numbers^44,45^.

### Supplementary Figures and Tables

**Figure S1.**
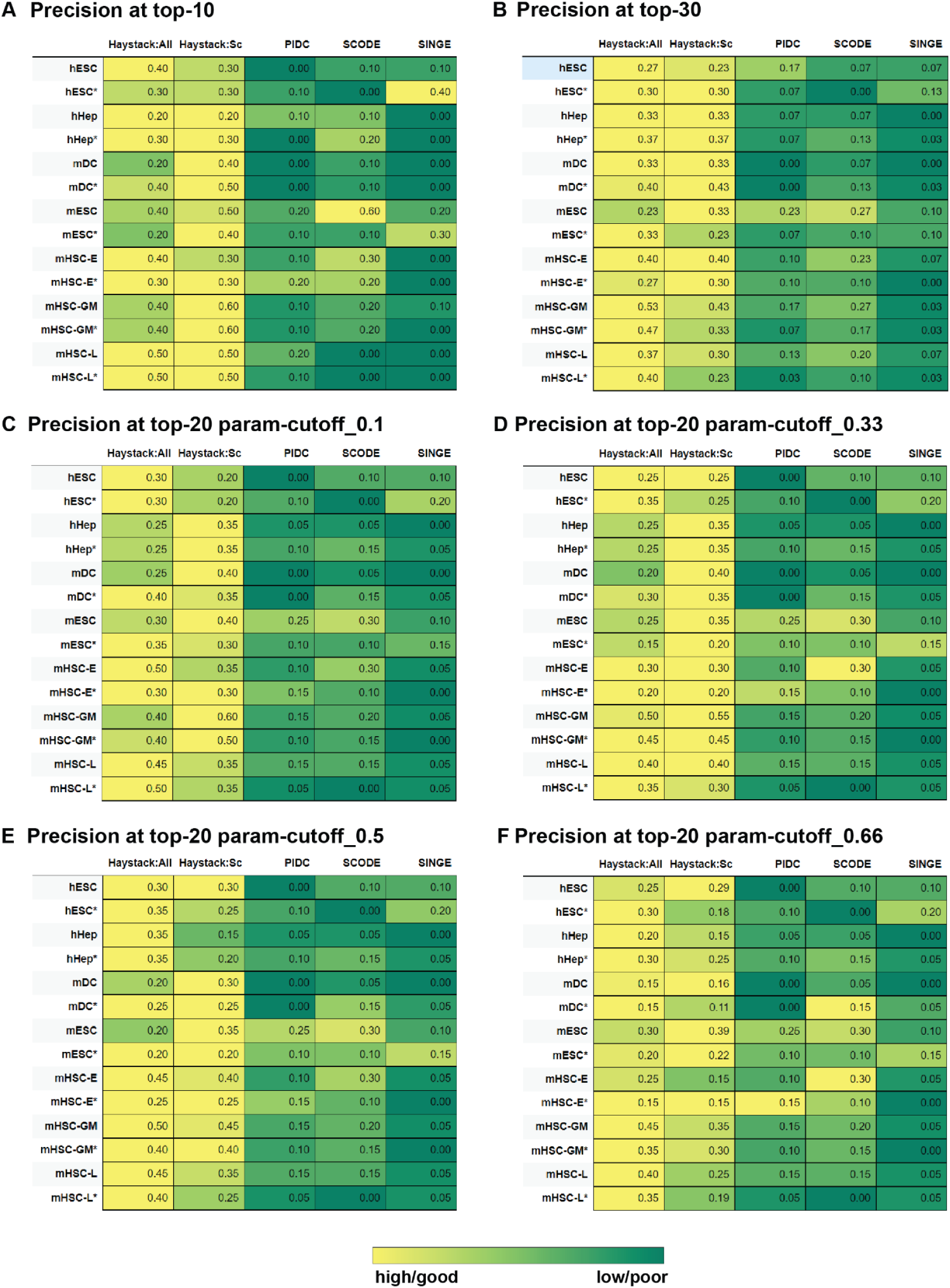
Benchmarking Haystack with other gene network inference methods. These plots accompany Fig 2 and show an extended comparison between Haystack and existing GRN methods. In all tabular plots, the yellow–green color gradient in each row is scaled to ensure a uniform maximum (yellow) across all rows. **A-B.** The tabular plot presents the precision of the top-10 (**a**) and top-30 (**b**) predictions of Haystack (All: source and target TFs, Sc: source-only TFs), PIDC, SCODE and SINGE on a variety of mouse and human cell types (Supplementary Note 2 for details on cell types). The ground-truth gene sets are sourced from ChIP-seq data, with an asterisk indicating evaluation against non-cell-type specific ChIP-seq. The precision results for top-20 predictions are shown in Fig 1h. **C-F**. For top-20 predictions, these tabular plots show the performance of Haystack over a variety of choices for the hyperparameter (“param-cutoff”) that controls the number of well-localized TFs selected after the initial optimal transport analysis: 0.1 (**c**), 0.33 (**d**), 0.5 (**e**), and 0.66 (**f**). For the results in Fig 1h, the parameter setting is 0.2.

**Figure S2.**
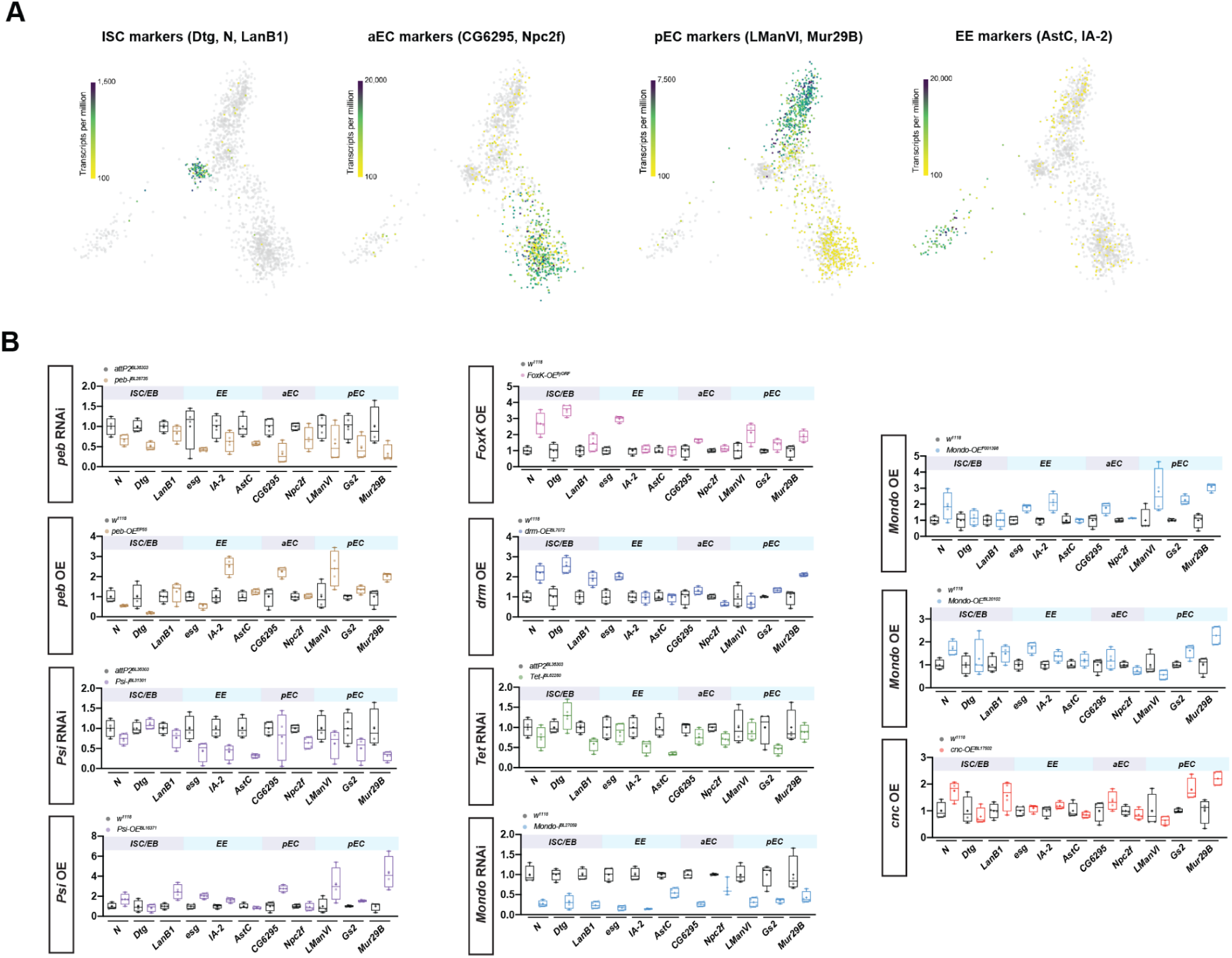
Analysis of the *Drosophila* midgut using qRT-PCR and confocal microscopy. **A.** UMAP plots representing the expression of intestinal cell marker genes for progenitors (*Dtg, N, LanB1*), aEC (*CG6295, Npc2f*), pEC (*LManVI, Mur29B*) or EE (*AstC, IA-2*) **B.** Bar graphs represent marker gene expression pertaining to the gut lineage validated by qRT-PCR of *Drosophila* gut mRNAs upon knockdown (KD) or overexpression (OE) of various source TFs such as *peb*, *Psi*, *FoxK*, *drm*, *tet*, *Mondo*, and *cnc*. The y-axis represents fold change compared to control normalized to *RpL32*.

**Figure S3.**
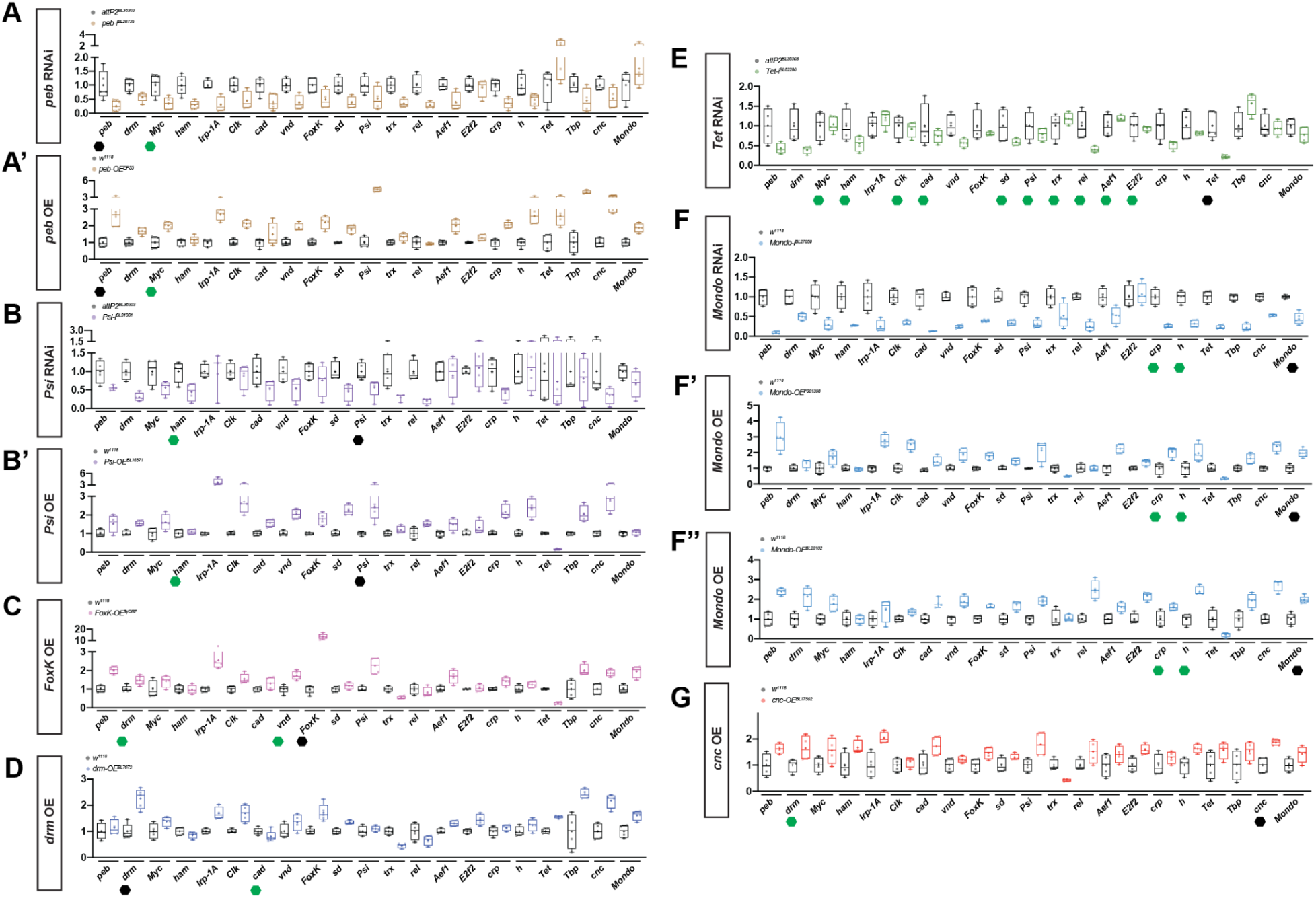
Marker gene expression and Source - Target validation by qRT-PCR. Bar graphs represent putative TF target gene expression pertaining to the gut lineage validated by qRT-PCR of *Drosophila* gut mRNAs upon knockdown (KD) or overexpression (OE) of various source TFs such as *peb* (A**, A’**), *Psi* (B**, B’**), *FoxK* (C), *drm* (D), *tet* (E), *Mondo* (F**-F’’**), and *cnc* (G). Panels are bar graphs representing the source TF (black hexagon) - target TF (green hexagon) validations by qRT-PCR of the aforementioned TFs. The y-axis represents fold change compared to control normalized to RpL32.

**Figure S4:**
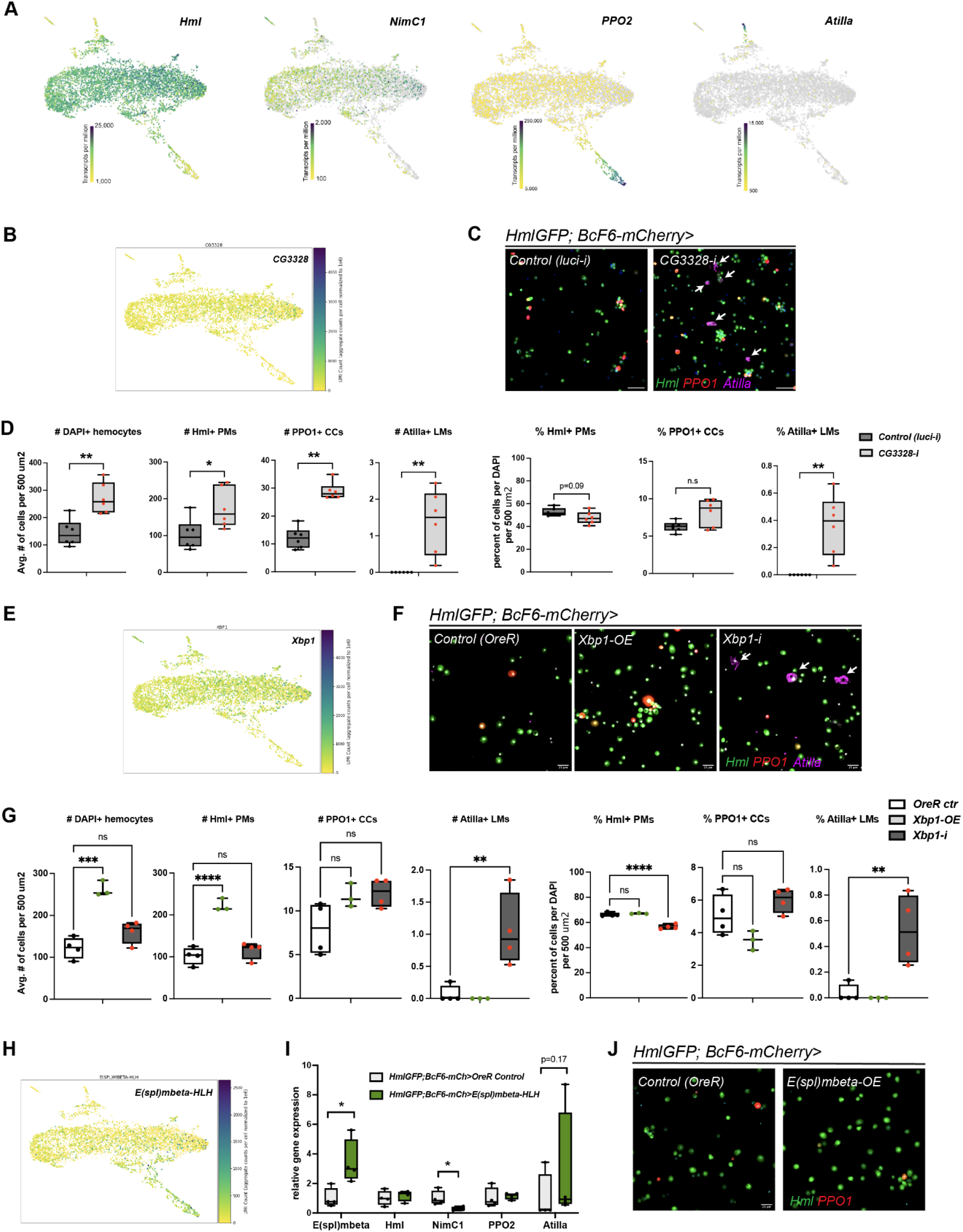
Haystack identifies known and novel TFs in *Drosophila* blood cell differentiation. **A.** UMAP plots representing the expression of blood cell marker genes *Hml*, *NimC1* (PM), *PPO2* (CC), and *Atilla* (LM). **B.** UMAP plot representing the expression of *CG3328*. C-D. Confocal images of *CG3328* knockdown (*CG3328-i*) shows production of Atilla+ LMs compared to luciferase RNAi (*luci-i*) controls (C, arrows). Bar graphs represent the cell counts which show an increase in the percentage of both PPO1+ CCs and Atilla+ LMs in *CG3328-i* (D). Note that total blood cell number (DAPI+ cells), Hml+ PMs, PPO1+, and Atilla+ LMs CCs are significantly increased upon *CG3328* knockdown. Non-parametric multiple t tests were used to calculate the statistics where * and ** represent p values <0.05 and 0.01, respectively. N=6 biological replicates. **E.** UMAP plot representing the expression of *Xbp1*. **F-G.** Confocal images (F) of *Xbp1*-overexpression (*Xbp1-OE*) and knockdown (*Xbp1-i*) shows increased blood cell number (in *Xbp1-OE*) and increased fraction of of Atilla+ LMs (arrows in **h**) compared to *OregonR* (*OreR*) controls (G). Note that *Xbp1-OE* causes an increase in the total blood cell numbers (DAPI+ cells) and Hml+ cells. One-way ANOVA was used to calculate the statistics where **, ***, and **** represent p values <0.01, 0.001, and 0.0001, respectively. N=3-4 biological replicates. **H-J.** UMAP plot representing the expression of *E(spl)mbeta-HLH* (H). qRT-PCR analysis shows that overexpression of *E(spl)mbeta-HLH* in blood cells decreases the expression of *NimC1* (I, N=4 biological replicates), validating its role in the PM^early^ → PM^late^ lineage. Confocal images of *OreR* control and *E(spl)mbeta-OE* show no marked changes in cell type composition (J). Non-parametric multiple t tests were used to calculate the statistics where * represents a p value <0.05.

**Figure S5:**
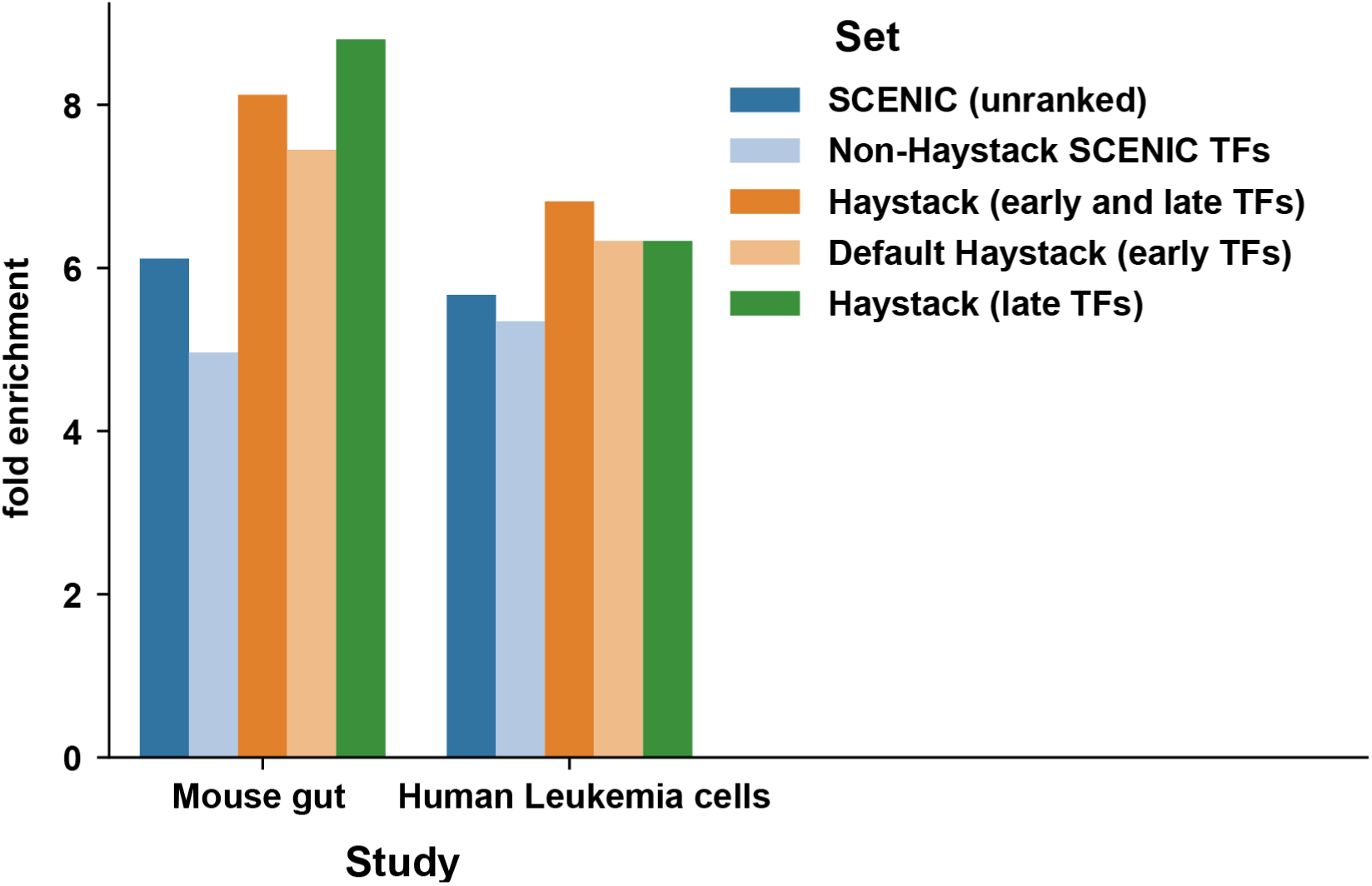
Additional evaluation of Haystack predictions using literature text mining metrics. This figure accompanies Fig 2H and shows the bar graph representing enrichment scores of TF predictions against ground-truth gene sets obtained by literature text mining. Here we show that Haystack performs well at recovering not just TFs active early in differentiation (“early TFs”) but also those that are active later in the terminal stages (“late TFs”). In all cases, Haystack’s TFs are selected from SCENIC regulons. As expected the regulons that Haystack does not select are not as informative as the ones it does select. Note that the method we used for our analysis in the main text focuses on the prioritization of early TFs (“Default Haystack”).

**Table S1.**
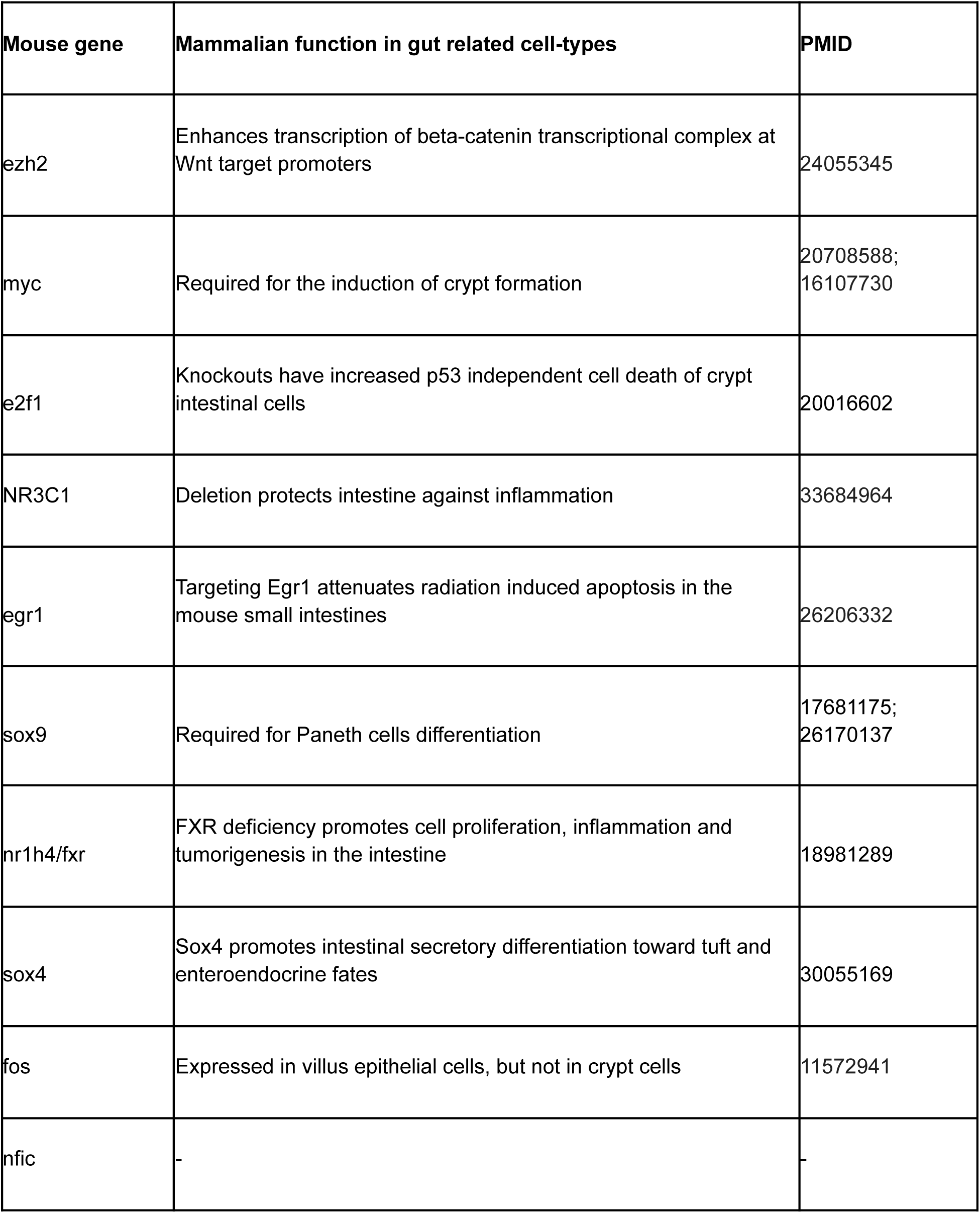
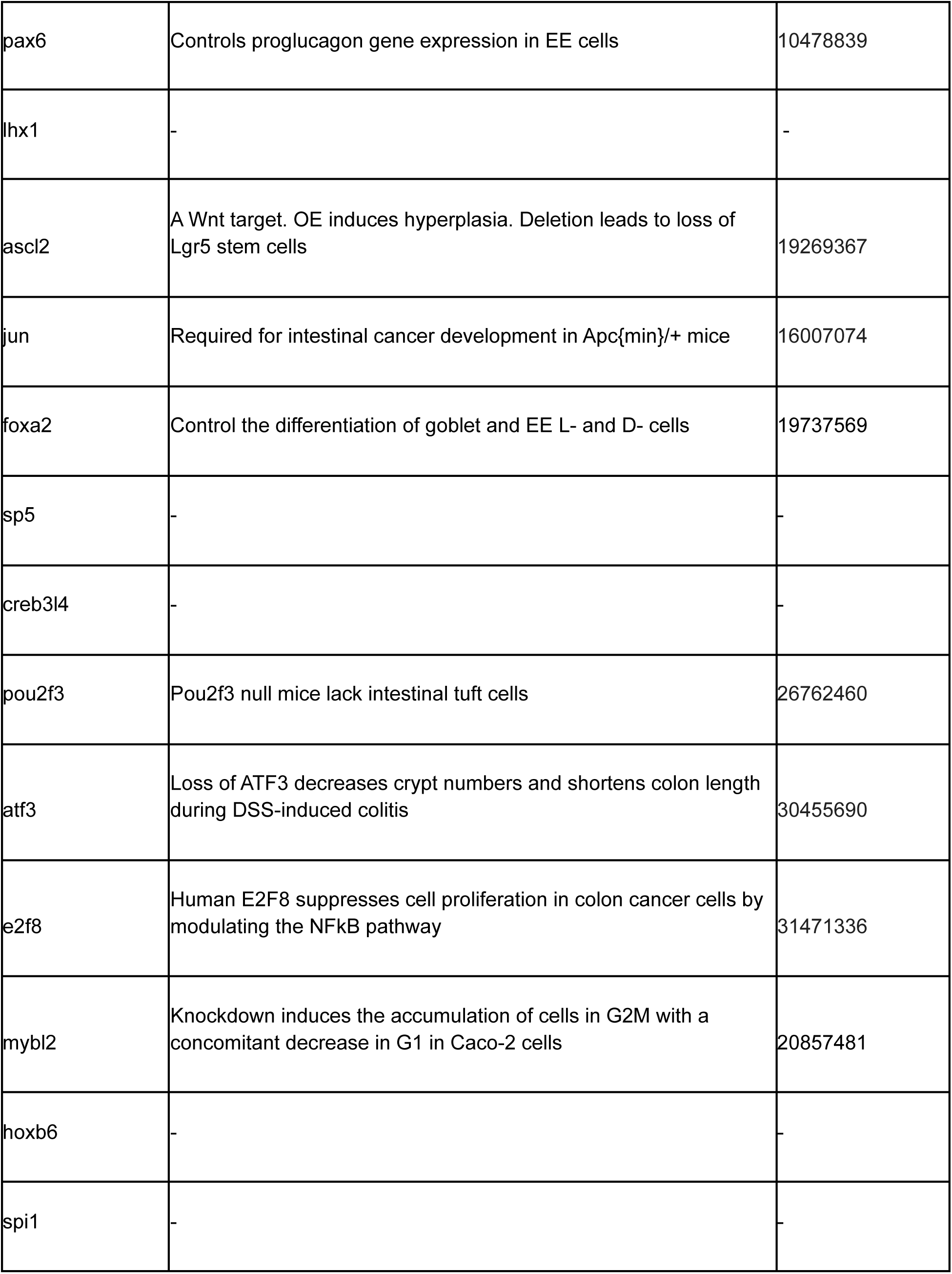
Haystack source TF predictions in the mouse gut

**Table S2.**
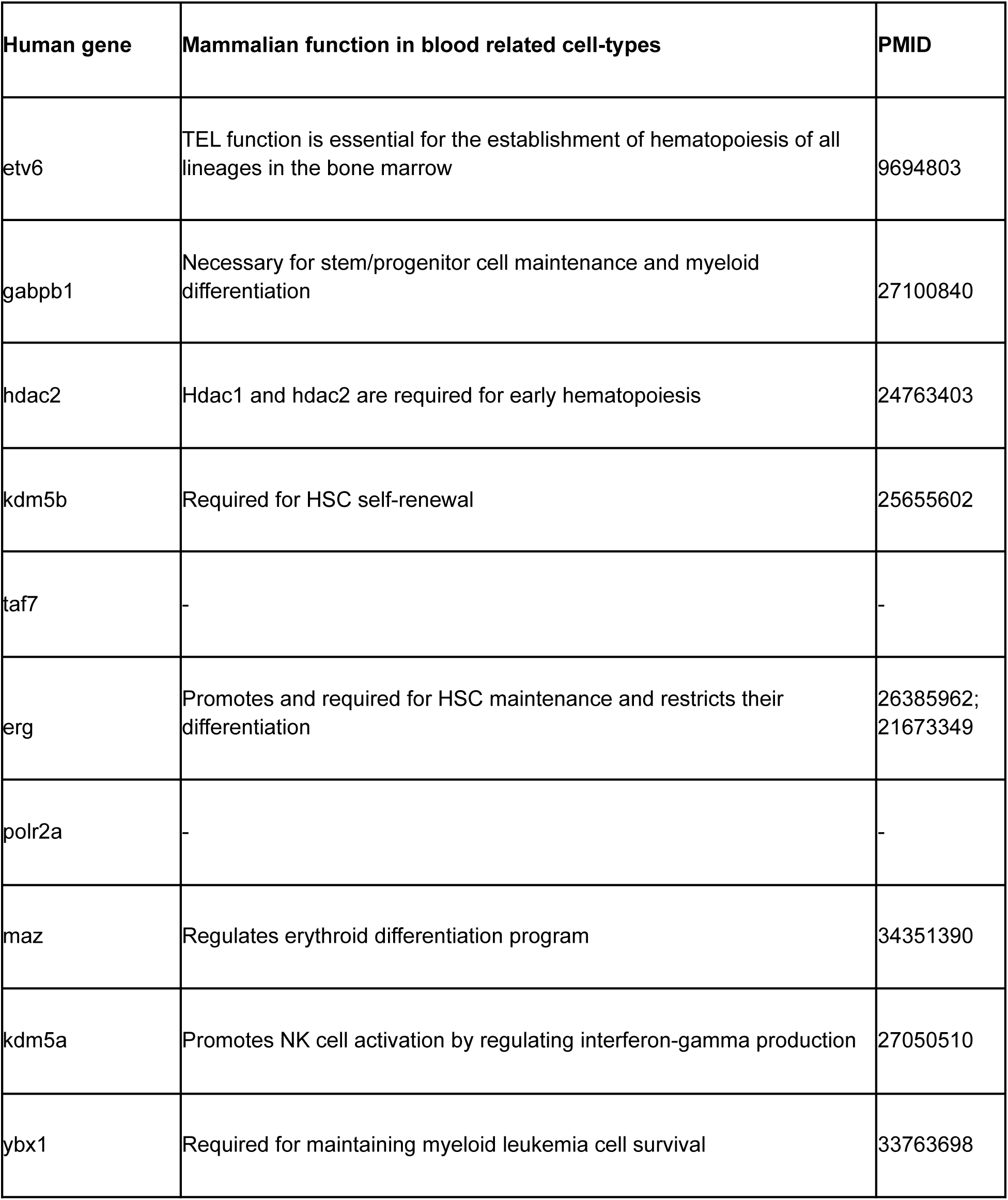
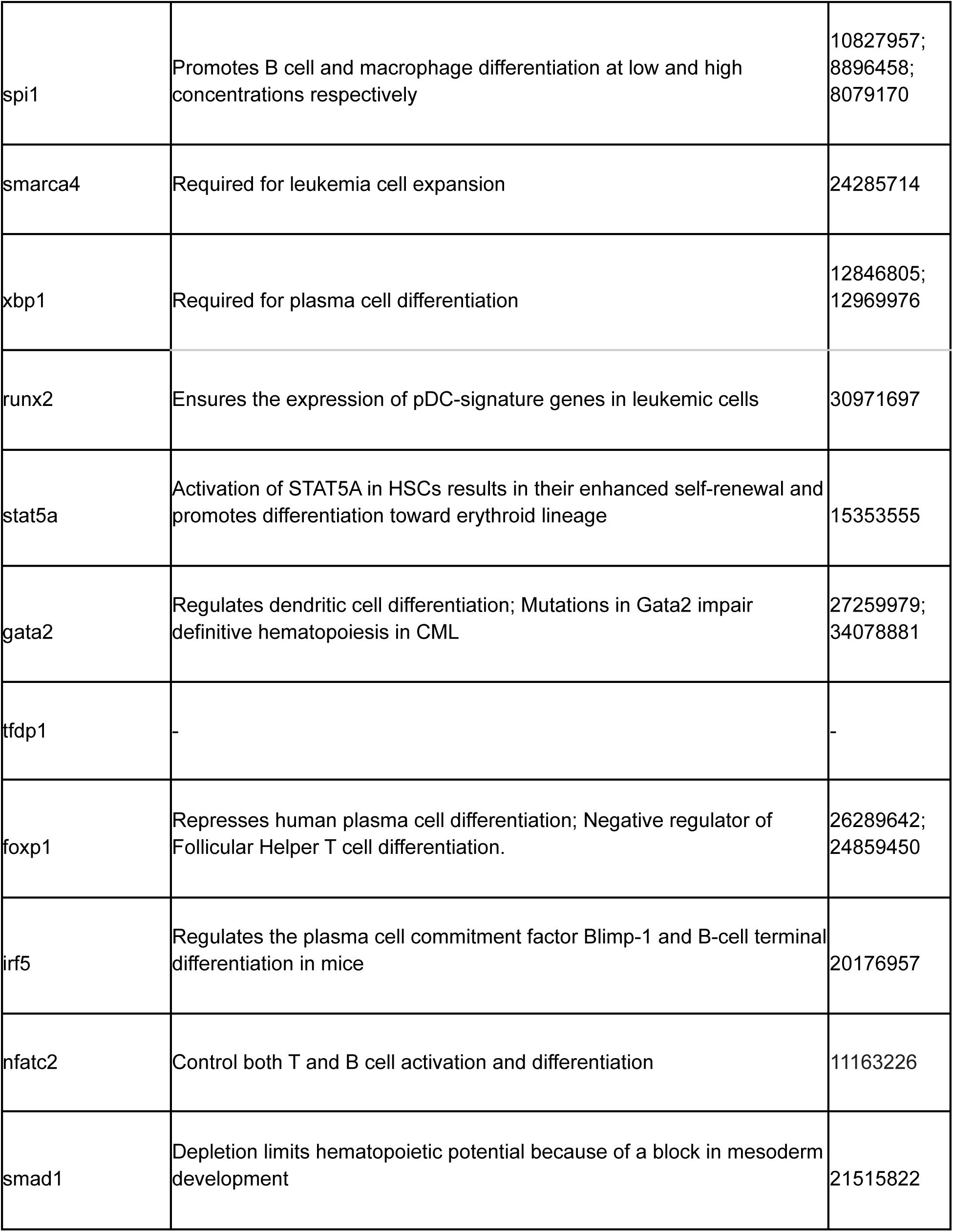
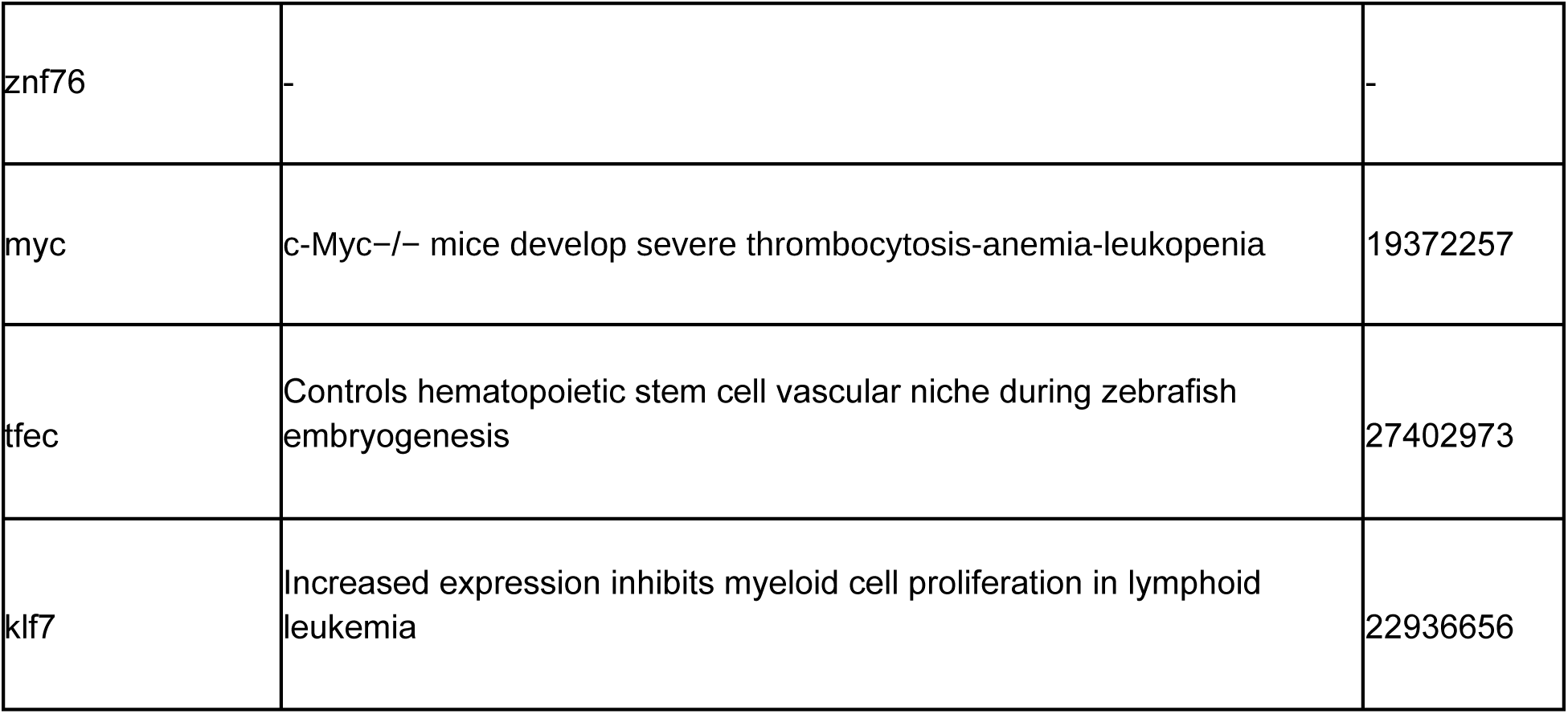
HACKSTACK source TF predictions in human leukemia

**Table S3.**
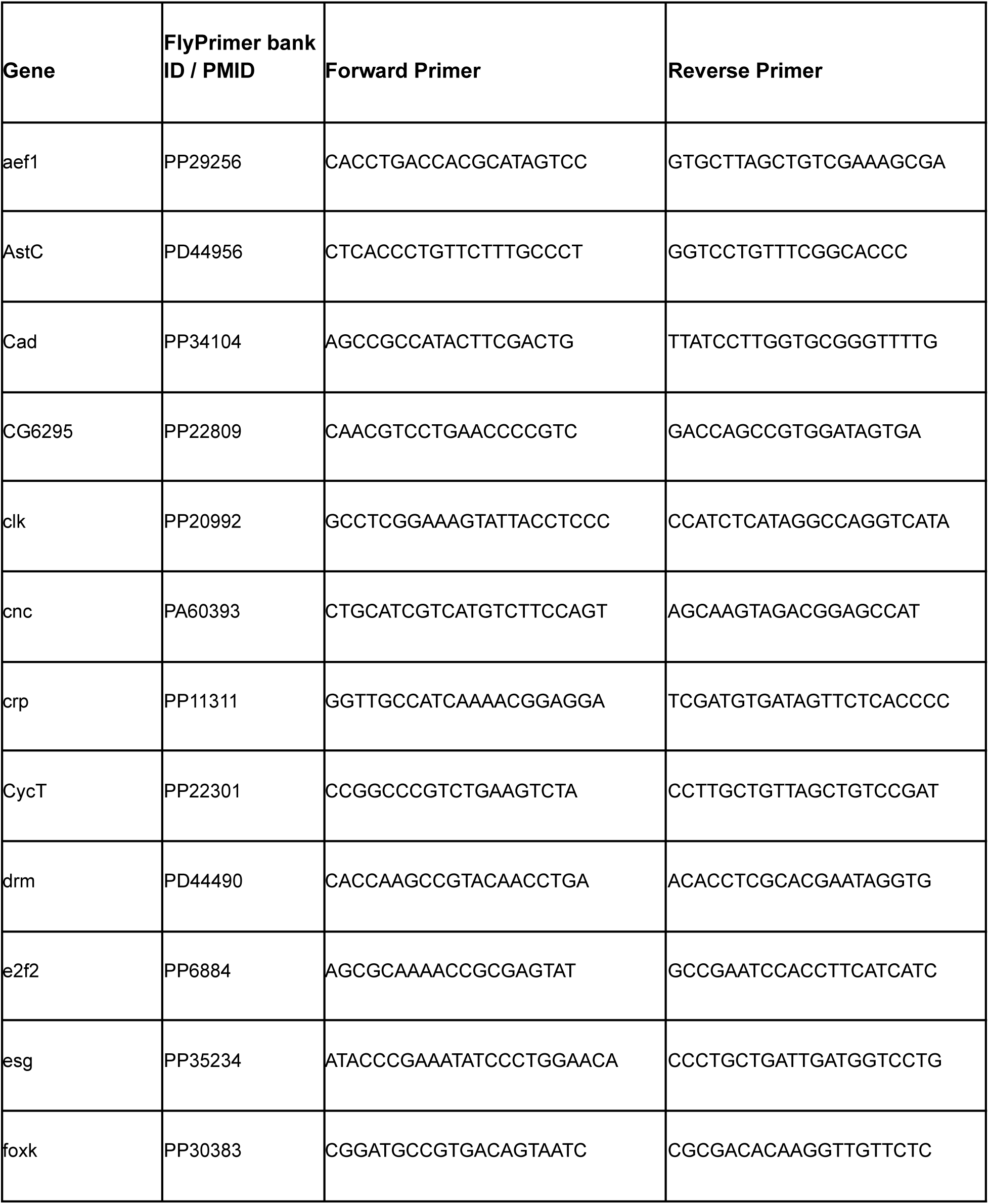
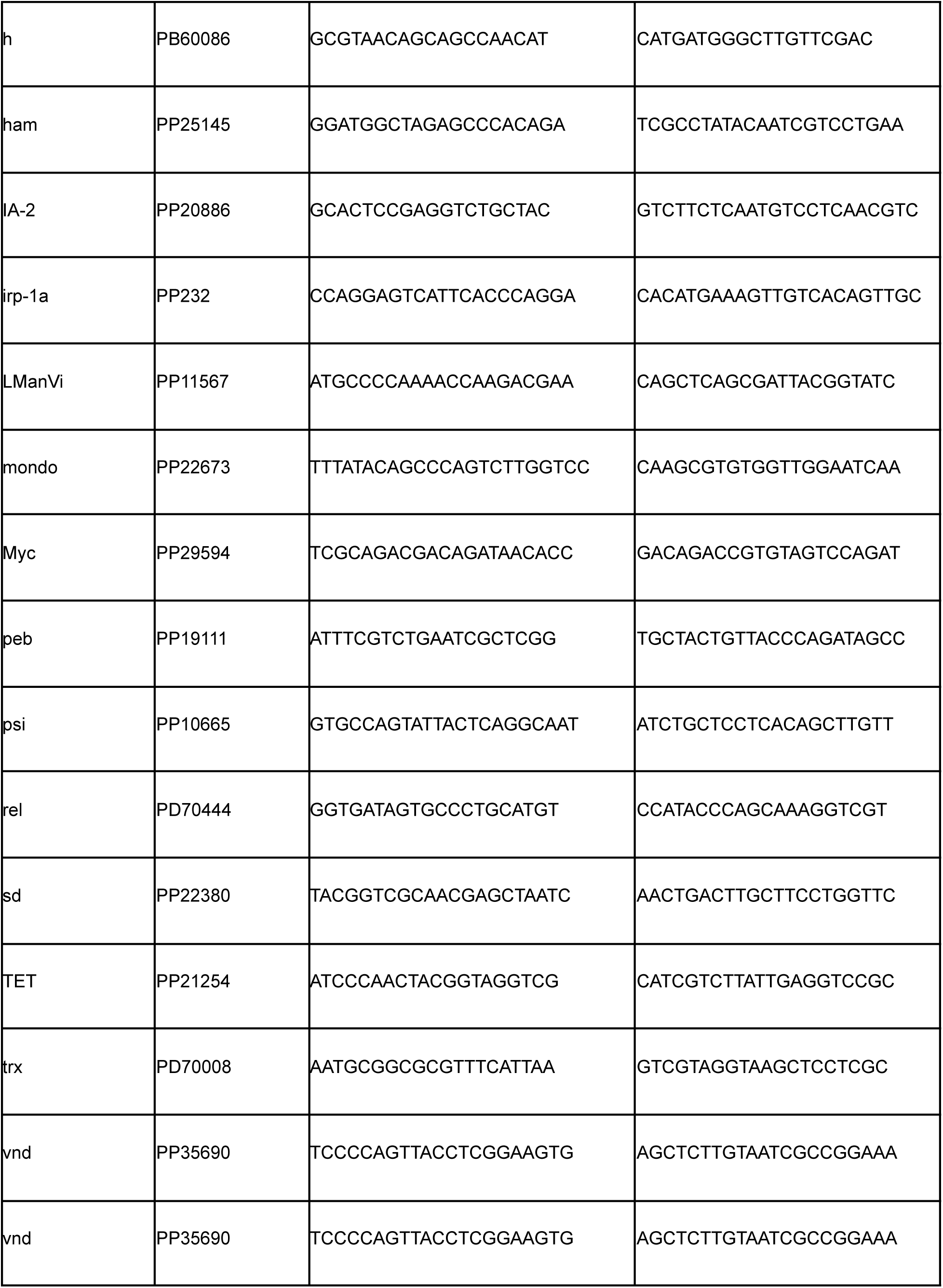
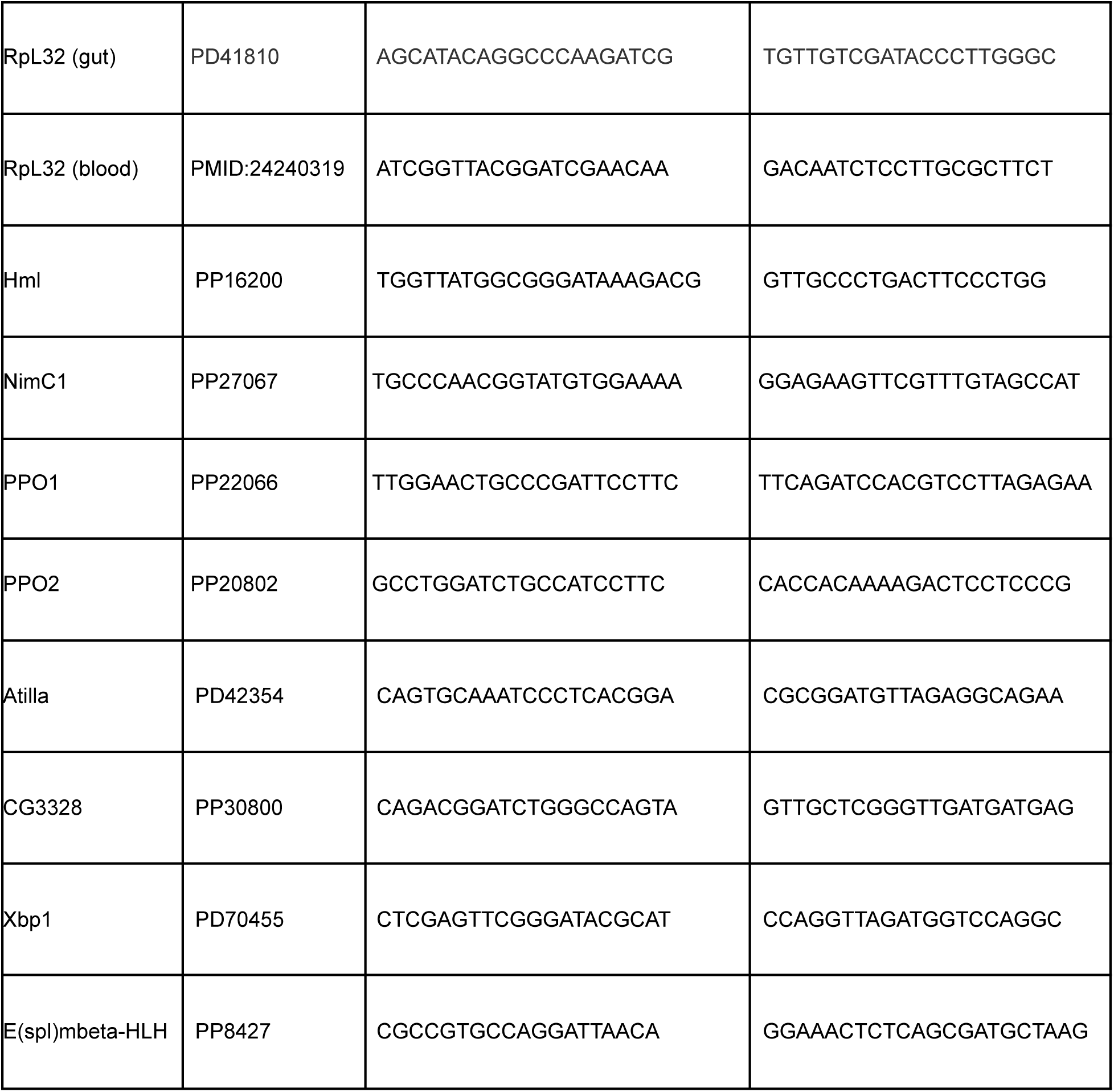
Primers used for qRT-PCR

